# Toward a comprehensive understanding of EEG and its analyses

**DOI:** 10.1101/2020.02.14.948968

**Authors:** A. F. Rocha

## Abstract

**Background:** EEG is the oldest tool for studying human cognitive function, but it is blamed to be useless because its poor spatial resolution despite it excellent temporal discrimination. Such comments arise from a reductionist point of view about the cerebral function. However, if the brain is assumed to be a distributed processing system recruiting different types of cells widely distribute over the cortex, then EEG temporal resolution and the many different tools available for its analysis, turn it the tool of choice to investigate human cognition.

**Proposal:** To better understand the different types of information encoded in the recorded cortical electrical activity, a clear model of the cortical structure and function of the different cortical column layers is presented and discussed. Taking this model into consideration, different available techniques for EEG analysis are discussed, under the assumption that tool combination is a necessity to provide a full comprehension of dynamics of the cortical activity supporting human cognition.

**Methodology:** The present approach combines many of the existing methods of analysis to extract better and richer information from the EEG, and proposes additional analysis to better characterize many of the EEG components identified by these different methods.

**Analysis:** Data on language understanding published elsewhere are used to illustrate how to use this combined and complex EEG analysis to disclose important details of cognitive cerebral dynamics, which reveal that cognitive neural circuits are scale free networks supporting entrainment of a large number of cortical column assemblies distributed all over the cortex.

**Conclusions:** Reasoning is then assumed to result from a well orchestrated large scale entrainment

## 1. Introduction

Electroencephalografy was born with the work of Hans Berger in the beginning of the XX^th^ century (Berger, 1929). It was the first technique to record, in a non-invasive way, the brain activity in humans. In his pioneer observations, Berger associated cognition to alpha waves and cerebral metabolism to cognition.

Ionic current in the cortex result from changes in neuronal and glial membrane conductance caused by intrinsic membrane processes and/or by synaptic actions. The net membrane current that results from such changes can be either a positive or a negative ionic current directed to the inside of the cell. Because the direction of the current is defined by the direction along which positive charges are transported, it may stated that at the level of the synapse there is a net positive inward current in the case of excitatory (EPSC) or negative one in the case of inhibitory IPSC currents. These currents are compensated by currents flowing in the surrounding medium since there is no accumulation of electrical charge in the extracellular space. Because cortical cells are distributed in a columnar topology in the cortex, a distributed passive source is created in the case of an EPSC and a distributed passive sink in the case of an IPSC. In this way a dipole configuration is created if the following is satisfied (e.g., Lopes da Silva, 2004; Olejniczak, 2006): 1) the cells are to be spatially organized with the dendrites aligned in parallel, forming palisades, and 2) the synaptic activation of the neuronal population are to occur in synchrony.

Thus, sources of the electrical recorded field are most probably compact regions of cortex (cortical column assemblies - CCA) whose local field activities at the cellular scale are similarly oriented by cortical geometry and partially synchronized at the cm^2^-scale.

The eletroecenphalogram (EEG) consists essentially of the summed electrical activity of the excitatory and inhibitory currents producing electrical and magnetic fields that may be recorded at a distance from the sources. These fields (*v* _*j*_ (*t*)) may be recorded by electrodes ***j*** at time ***t*** from sources at short distance - called local field potentials, or from the cortical surface or even from the scalp - called EEG in the most common sense. Lopes da Silva, 2004). Electroencephalografy gives us the possibility of studying brain functions with a high time resolution, although with a relatively modest spatial resolution.

One of the key issues in electroencephalography is to identify the cortical sources generating the recorded EEG. This is the so called inverse problem. There is no unique solution for this question because *v* _*j*_ (*t*) measurements do not contain enough information about their generators or because these generators are many. This gives rise to what is known as the non-uniqueness of the inverse solution. In this way, realistic source analysis of EEG potentials requires objective biophysical models that incorporate the exact positions of the sensors as well as the properties of head and brain anatomy, such that appropriate inverse techniques can be applied to map surface potentials to cortical sources. Hence, it can be stated that a perfect tomography can not exist and many solutions have being proposed (see for discussion Grech, et al, 2008 and Pascual-Marqui, R. D, 1999). Among such techniques, LORETA has been very popular and provides an acceptable solution, besides being freely available at Internet. Although precision of EEG tomography is very dependent on the density and localization of the electrodes (Song, et al, 2015). an acceptable solution may be achieved with use of 20 electrodes (Michel et al, 2004).

In the early years, leading-edge electroencephalographers often designed and constructed their own apparatus or alternatively, acquired and adapted equipments intended for other uses. Many of these electroencephalographers have background into communications, audio engineering, office machines, etc (*Collura, 1993*). As commercial systems became available and evolved considerably, researchers continued to search to add extra capability to these new devices and to find new ways to improve signal analysis. EEG analysis remained strongly visual until recorded electric activity could be digitized and stored for further processing taking advantage of the continuous and enormous development of computers. Quantitative EEG (qEEG) analysis was then born (e.g., John, et al, 1988 Nuwer, 1997).

Fast Fourier Analysis is one of the most popular technique on qEEG. It decomposes the recorded EEG wave into a series of sinusoidal waves called EEG band frequencies. Normally up to 6 bands are considered: delta band characterized by frequencies smaller than 4 Hz; theta band characterized by frequencies between 4 to 7 Hz; alpha band composed by frequencies between 8 and 13 Hz; beta band with frequencies between 15 and 30 Hz and gamma band characterized by frequencies greater than 30 Hz. The power spectrum of each band frequency provides a measure of its contribution to the recorded EEG. Many different cognitive functions have being associated to these band frequencies (e.g., Aftanas etl, 2001; Christof et al, 2016; Doesburg. Kitajo and Ward, 2005; Klimesch, 1999), Klimesch, 2012; Piai ET AL, 2015 and Weisset al, 2000).

Another frequently used technique to study EEG activity associated to cognition is called Evoked Related Potential (ERP) or Activity (ERA) and imply to average the recorded EEG following a particular sensory stimulation or the onset of a given cognitive activity such as speech understanding; picture recognition, etc. Many distinct ERA components are identified by its polarity and time occurrence, eg: N50 - negative wave peaking at 50 ms after stimulus onset; P100 - positive wave peaking with a latency of 300 ms; N400 - negative wave peaking at 400 ms; P600 - positive wave peaking at 600 ms, etc. (e.g., Donchin, et al 1975; Kutas, and Hillyard, 1980; Morgan; Hansen and Hillyard, 1996 and Picton, et al 1974 *and* Regan,1966).

Using ERA traditional analysis, researchers 1) average a set of EEG data trials or epochs time-locked to some class of events, obtaining an EEG waveform (ERP) or 2) study the average changes in the frequency power spectrum of the whole EEG data time locked to the chosen events, producing a two-dimensional image that we call the event related spectral perturbation (ERSP). But even together, the ERP and ERSP do not fully capture the mean event-related dynamics expressed in the data. (Makeig et al, 2004).

A reductionist approach of cortical function proposing that complex cognitive functions may be assigned to specific events and cortical places, contrasts to a more recent proposals taking cerebral dynamics to be that of a distributed processing system recruiting wide sets of CCA located at many different cortical areas to take charge of the distinct tasks of a complex cognitive reasoning (e.g., Baars, 1988, 1997, Baars et al, 2013; Iturria-et al, 2008; Hipp et al, 2012; Raffaele et al, 2008; Rocha, 199; Rocha, et al, 2004, 2011, 2017 and Smit, et al, 2008). EEG high temporal discrimination is very adequate to sample brain activity associated to this distributed processing, and the actual available tools for EEG analysis, if used in combination, certainly are of great help to study details of the parallel, concurrent and massive columnar activity supporting human reasoning. We have been successfully combining some of these tools to study different types cognitive tasks (Rocha, et al,; 2013; 2014; 2015, 2017 and Vieito, Rocha and Rocha, 2913).

The purpose of this paper is to discuss how and why a combined use ERA, LORETA source identification and other techniques provides a better understanding of the cognitive cerebral dynamics. Unpublished data from our investigation on language understanding (Rocha, Foz and Pereira Jr., 2015) will be used to illustrate this combined use of tools. In this study, a small text introduces a quiz about fruits, tools or professions. After the listening or reading of the quiz, five pictures about fruits, tools or professions are proposed as solution and volunteer has to select one as the correct answer to the quiz. Three EEG epochs were selected for analysis: EEG epoch Q - 2 seconds from the beginning of text presentation; EEG epoch S - 2 seconds from the beginning of pictures display and EEG epoch C - 2 seconds prior choice selection.

The paper is organized as follows:

1. section 2.1 briefly presents and discuss the columnar cortical structure and the different types of electrical activity that may be recorded at the different columnar layers;
2. section 2.2 correlates the recorded *v* _*j*_ (*t*) to columnar cortical activity and to the neural net topology given rise to a entrained cortical activity modulated by intrinsic and extrinsic columnar circuits;
3. section 2.3 defines and formalizes a measure (*E*(*j*)) of this cortical activity entrainment recorded by each electrode ***j*** based on EEG correlation analysis;
4. section 3.1 shows how to use Factorial Analysis to disclose *E*(*j*) patterns,
5. section 3.2 shows how to use sinusoidal segmentation functions to identify Event Related Activity (ERA) components;
6. section 3.3 discusses the Oscillatory Cortical Activity that may be identified by using Fast Fourier techniques;
7. section 3.4 shows how Factor Analysis are to be used in detecting *E*(*j*) covariation patterns that reveals a complex concurrent entrainment of the different neural circuits involved in reasoning;
8. section 3.5 shows how to use LORETAs to solve the inverse problem of localizing ERA sources;
9. section 3.6 discusses the network properties of cortical columnar activation;
10. section 3.7 shows how to use LORETAs to solve the inverse problem of localizing cortical oscillatory sources;
11. section 3.8 correlates LORETAs and FA analysis results and shows how this tool combination provide final details of the columnar cortical entrainment dynamics, and
12. section 3.9 proposes a regression model for reversal engineering EEG reconstruction from the identified cyclic ERA activity and provides some concluding remarks

## 2. Theoretical background

### 2.1 Cortical structure and the genesis of electrical currents

Korbian Brodmann divided the cortex into 52 different areas (Figure 1) concerning their cito-architectonic characteristics (Brodmann, 1909). The columnar functional organization of the cells (Figure 1) in these different areas was discovered by Mountcastle, 1957 and confirmed by Hubel, Wiesel and Wiesel, 1977.

**Figure 1.**
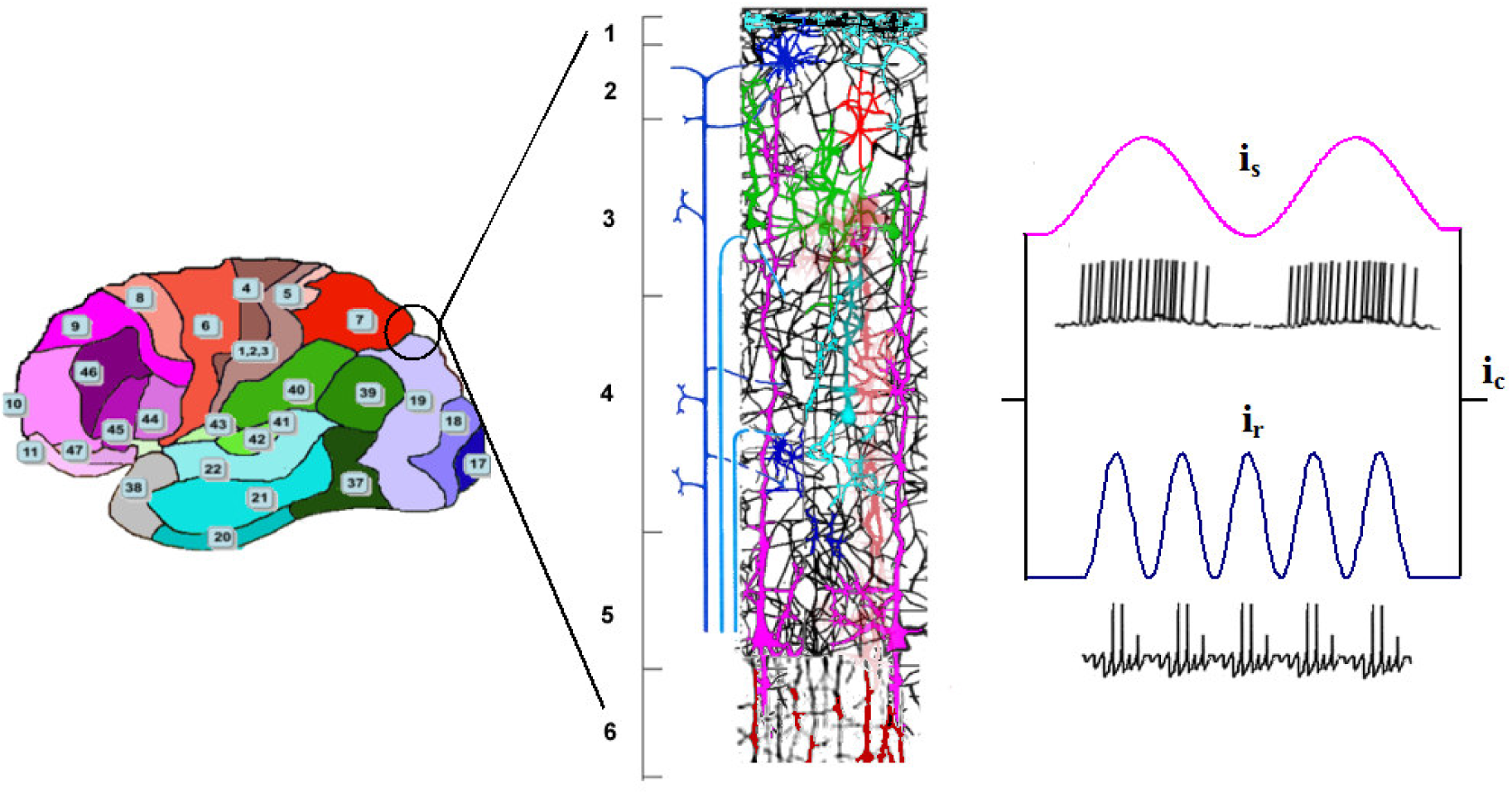
Brodmann areas and the structure of a cortical column composed by 6 layers characteristics of the Eulaminated II cortex (Barbas, 2015). Although columnar organization varies for different cortical areas, the structure shown in this figure will used, for illustration purpose, in the present paper. Tonic and Burst oscillatory columnar electrical activity is due to both intrinsic membrane properties of rhythmic cells or inhibitory circuits involving interneurons from the same or neighboring columns. Due to the parallel columnar organization, the oscillatory activity at the distinct layers (*i*_*s*_ (*t*),*i*_*r*_ (*t*), etc.) sum up to generate a unique columnar electrical current (*i*_*c*_ (*t*)).

Different neural systems (serotoninergic (5-HT), monoaminergic (NA), colinergic (Ach) and histaminergic (HA)) modulate neural activity of columnar cells (Figure 2) that may vary from burst to tonic oscillatory activity (e.g., Rocha,. Pereira Jr and. Coutinho; 2001).

**Figure 2.**
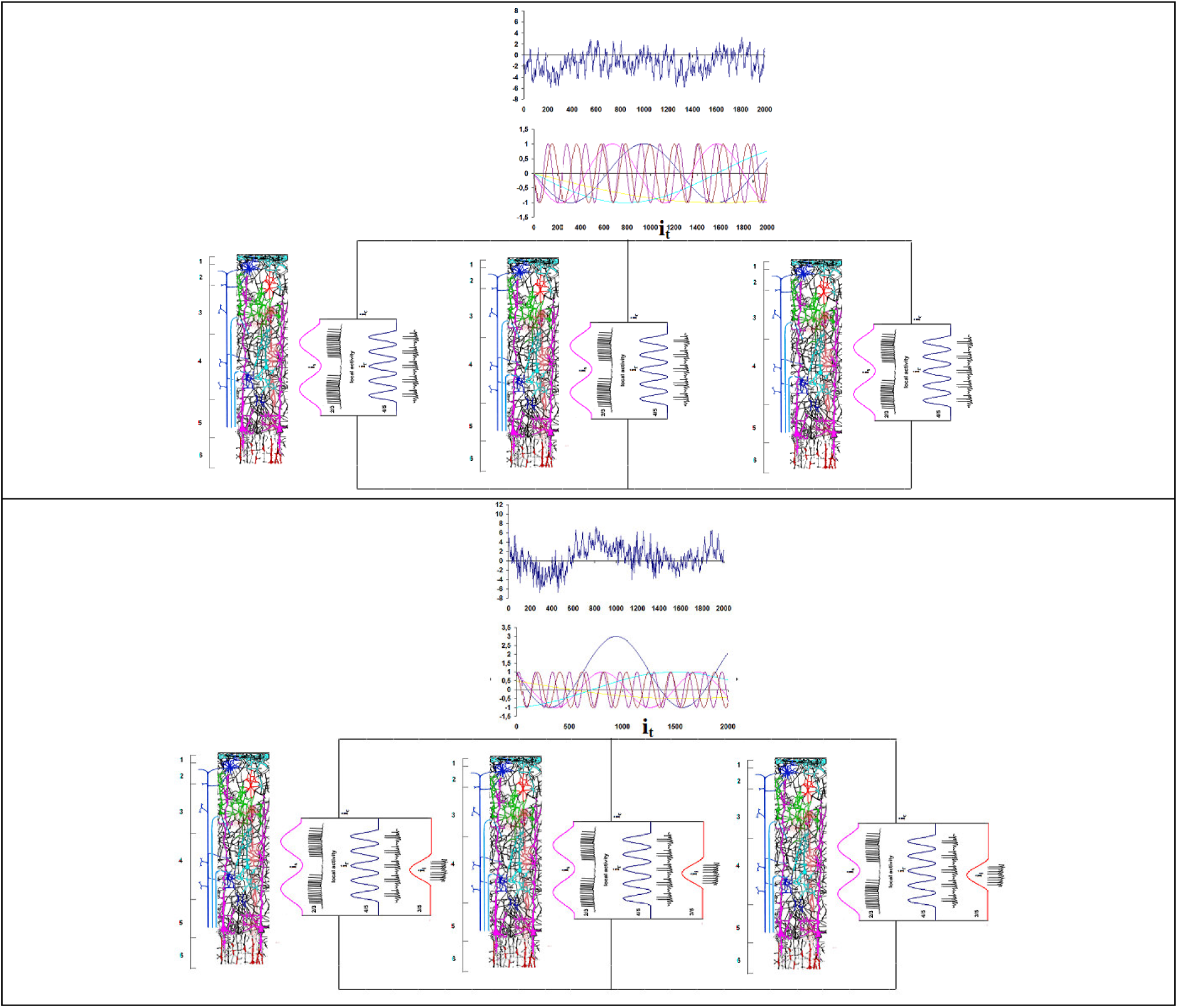
Columnar activity - Above: If there is no activity at the input fibers (resting activity), local oscillatory activity is not entrained and i_c_ tends to be low and noisy. Below: Activity of input fibers organize intrinsic columnar oscillatory activity and add an input current *i*_*i*_ (*t*) to the total columnar ionic current *i*_*c*_ (*t*). The entrained activity promoted by *i*_*i*_ (*t*) creates a larger *i*_*c*_ (*t*) in contrast to the low resting *i*_*c*_ (*t*). In addition, input fiber activity entrains the activity of the neighboring columns too, whose currents sum up generating a current *i*_*t*_ (*t*) representing the electrical activity of an entrained columnar assembly (CCA).

Astrocytes make contacts to both pre-synaptic terminals and postsynaptic dendrites and are influential upon neuronal activity, too (e.g., Pereira and Furlan, 2010).

Oscillatory activity at the different layers (Figure 2) may be dependent on their membrane properties or due to local neural circuits involving cells of the same or neighboring columns (Contreras and Llinás, 2001; Moritz et al, 2009; Shuzo, and Harris, 2009). Input activity from thalamus is another important influence over columnar electrical activity, mostly the oscillatory activity originating on the reticular thalamic neurons (McCormick, et al, 2015; Olejniczak, 2006; Rocha, Pereira Jr. and Coutinho2 2001 and Steriade, McCormic and, Sejnowski, 1993). Finally, input from pyramidal neurons from other nearby or distant columns as well as from parallel fibers originating in neighbor columnar interneurons complete set of determinants of the columnar electrical activity (e.g., Rocha,. Pereira Jr and. Coutinho; 2001)

Many different ionic currents are generated by cellular columnar activity at the different layers (Figures 1) having properties that are dependent on the electrical activity in the input fibers (Burgess, 2012, 2013; Gilley and Sharma, 2010; Giraud and Poeppel, 2012 and Shuzo and Harris, 2009). At low and asynchronous input activity, local oscillatory activity at the distinct layers tend to be uncoupled. Due to the columnar cellular organization these distinct layer activity create different currents (*i*_*s*_ (*t*) and *i*_*r*_ (*t*)) that sum up generating a total columnar current *i*_*c*_ (*t*) that is low and noisy (Figure 2). Increase of the input activity triggered by sensory stimulation or internal activity created by reasoning, entrains not only the intrinsic activity of each column but also the activity of a defined number of neighbor columns or a Cortical Columnar Assembly (CAA). This entrainment increases the value of the different *i*_*c*_ (*t*) that sum up into a larger current *i*_*t*_ (*t*) whose value heavily depends on the number of the entrained columns or CAA size (Figure 2).

### 2.2 The recording of the electrical columnar activity: The Encephalogram

As discussed above, the value *i*_*i*_ (*t*) current at time ***t*** is the summation of *i*_*c*_ (*t*) generated by the entrainment of ***n*** columns that is

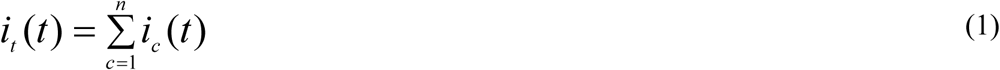

This current creates the electrical field

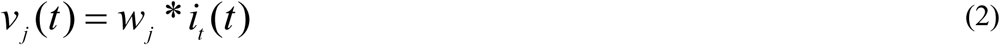

under each electrode *e* _*j*_ outside the skull depending on the electrical resistance *w*_*j*_ between the electrode and the CAA site. The value of ***n***, as pointed before, is dependent on the value and type of the input current *i*_*i*_ (*t*), such that

1. ***n*** decreases as *i*_*i*_ (*t*) decreases and becomes asynchronous and
2. ***n*** decreases as *i*_*i*_ (*t*) increases and its synchrony enhances.

In such a conditions, the value and evolution of *v* _*j*_ (*t*) recorded by the electrode *e*_*j*_ is mostly determined by the behavior of *i*_*i*_ (*t*),

Many different CCAs are continuously being organized at the different cortical areas by the incoming input activity, such that each electrode makes a distinct 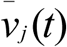 record of the different electrical fields that *v*_*j, d*_ (*t*) generated by these CCAs because the distance between each electrode *e*_*j*_ and the ***k*** different CCAs (Figure 5) are specific to each *e*_*j*_, that is

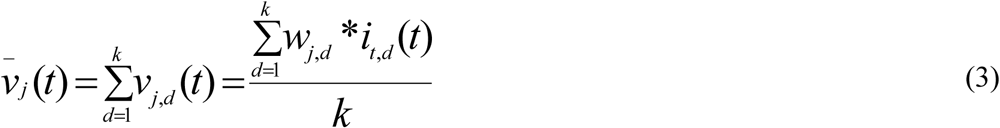

**Figure 3.**
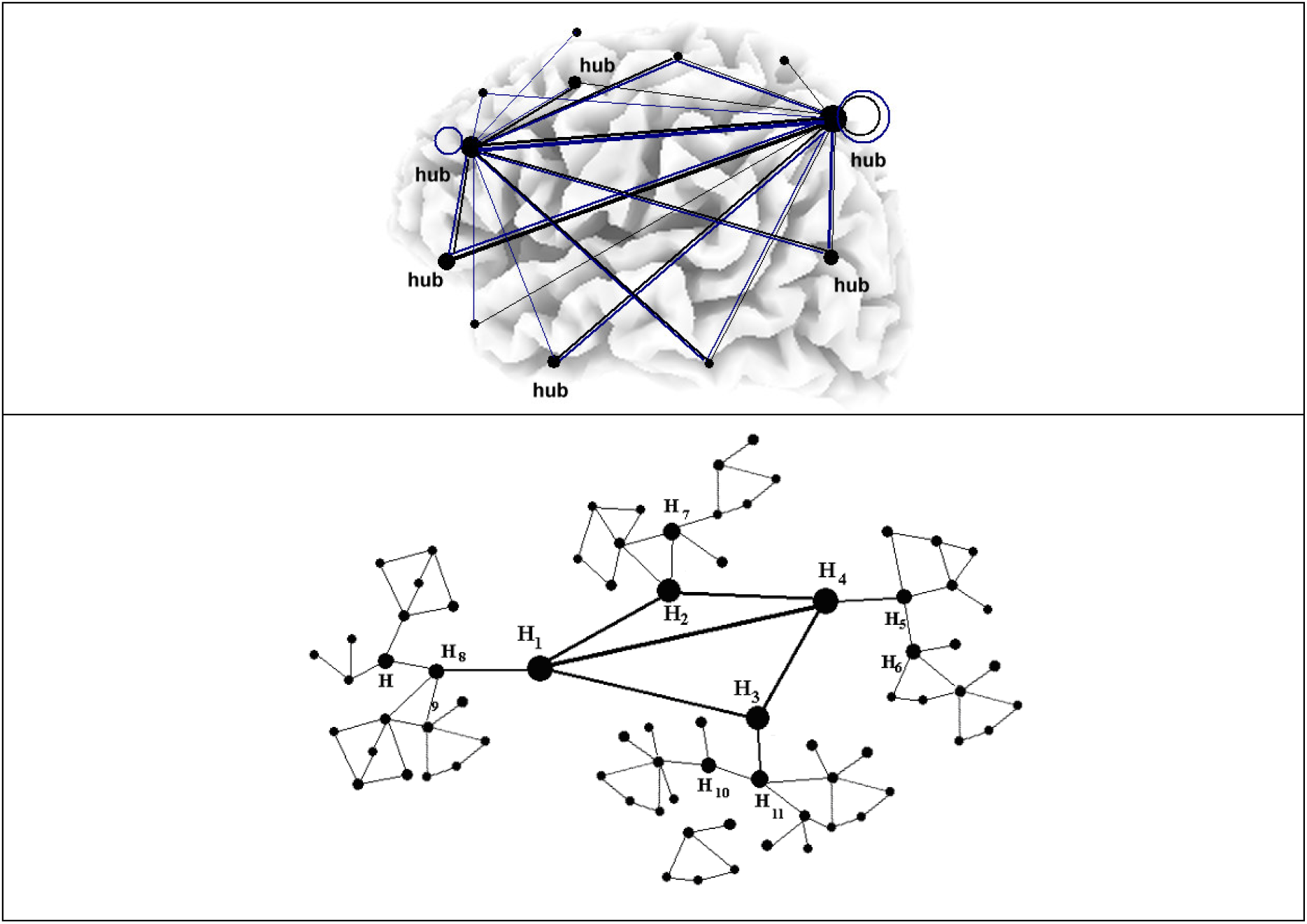
The scale-free topology for message exchange in cortex. Node hubs in these networks efficiently exchange information and distribute it to subordinate nodes

**Figure 4.**
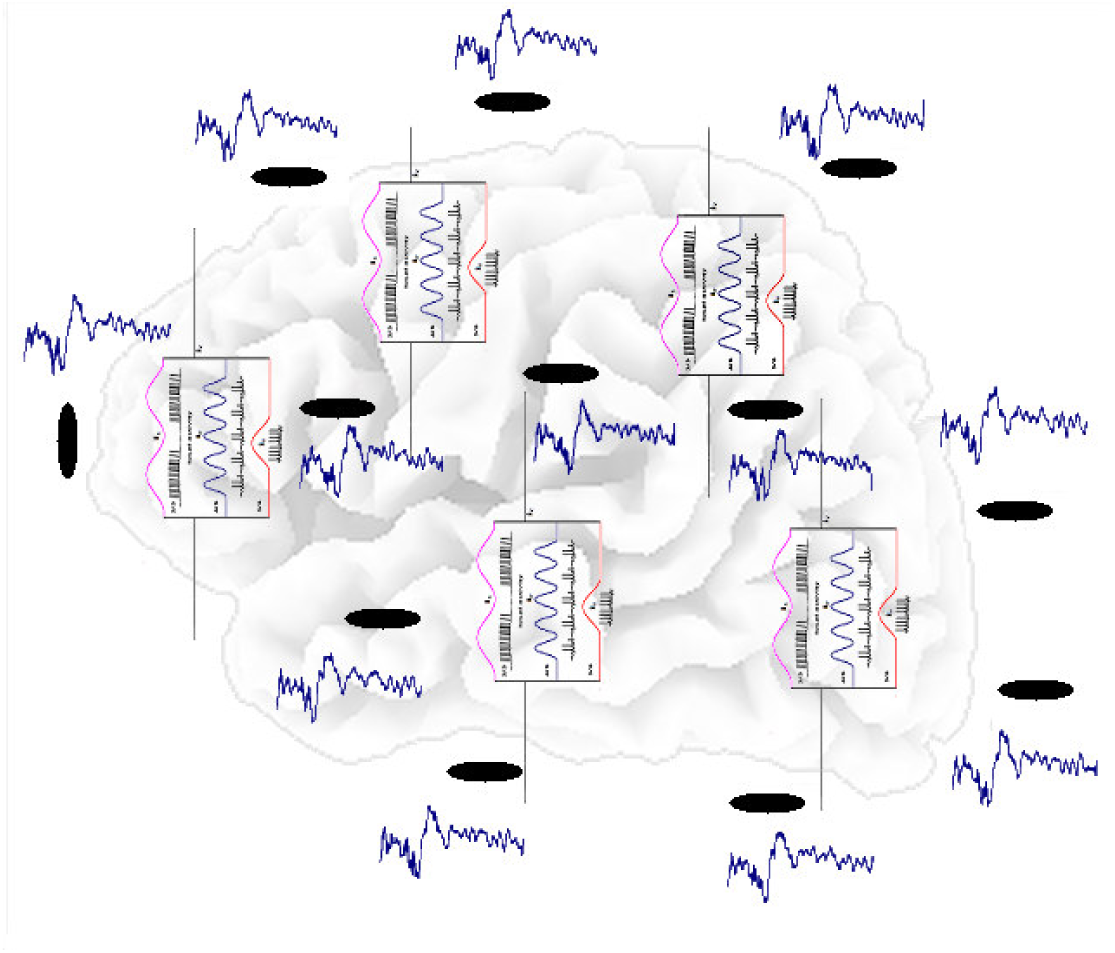
Scale free topology favors the entrainment of the activity recorded by the distinct electrodes. The figure shows the averages of the recorded activity EEG epoch Q calculated for 41 volunteers and each recording electrode. Similar results were obtained in studies of other different reasoning processes ((e.g., Rocha et al, 2017)

**Figure 5.**
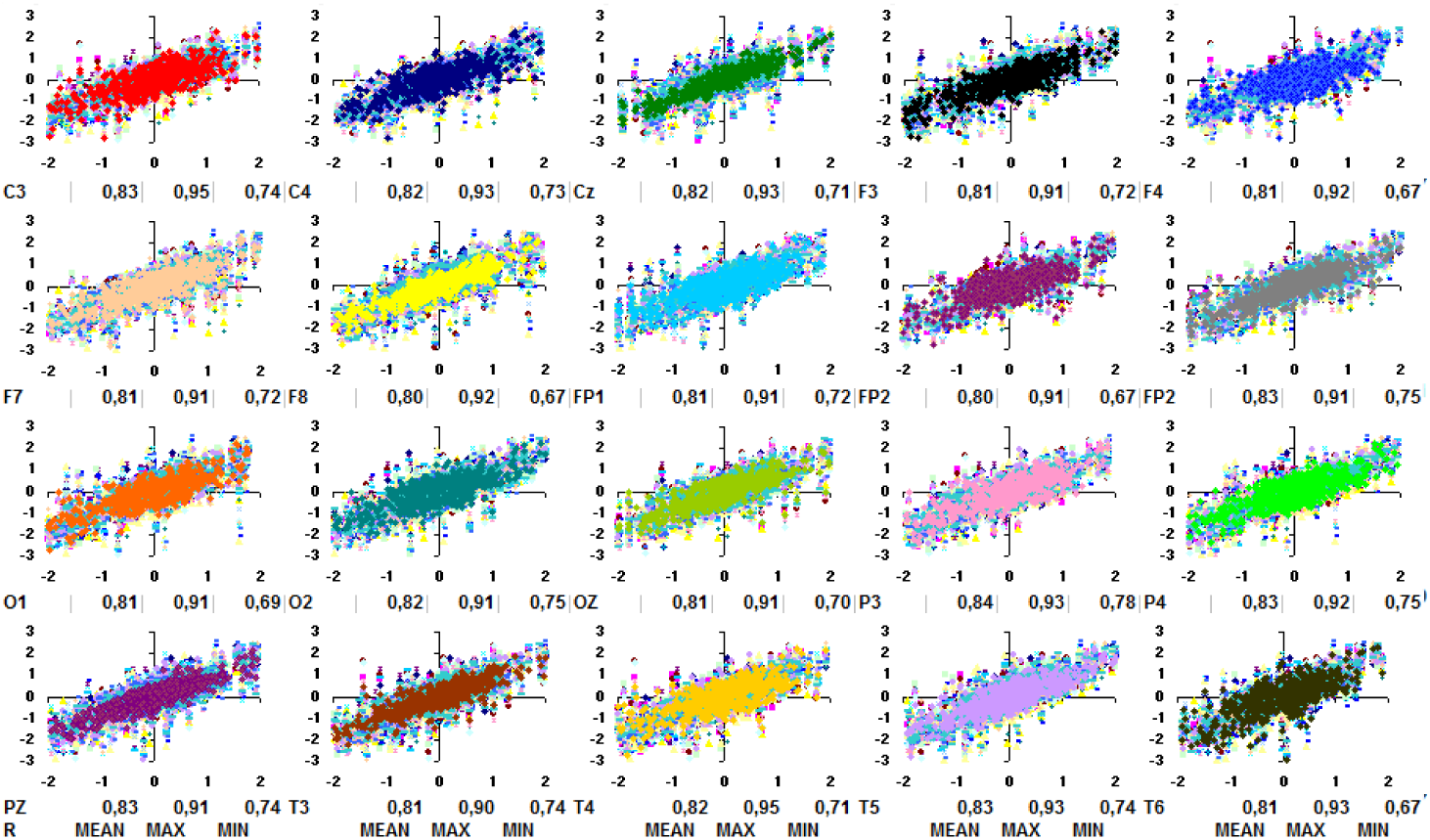
Correlation between EEG epoch Q activity recorded by the distinct electrodes. Mean - average of the regression coefficient R obtained for correlation between the selected electrode and the remaining 19 ones. Max and Min - maximum and minimum calculated R. Similar results were obtained in studies of other different reasoning processes (e.g., Rocha et al, 2017)

There is a arcane debate if ERA patterns are signals super-imposed upon and independent on the ongoing *v* _*j*_ (*t*), or if they results from phase alignment of the ongoing columnar oscillatory activity, or some combination of the two (Burgess, 2012). Here, ERA is proposed to result from this later combined process. Because of this, they cannot be explained by specific activation of particular set of neurons, but have to be understood mostly as resulting from an entrained oscillatory activity at multiple cortex locations (Burges, 2012; Klimesch et al,2007).

To reduce the costs of message spread in the brain and to support an entrained oscillatory activity, nature developed some kind of special networks, named scale free networks (Figure 3) that are very efficient in spreading information among their nodes, using some preferential interconnection between some special nodes (CCA) that function as hubs for message spread. Efficient networks have a low number of highly connected hubs that are heavily connected between them and to the local subordinate CCAs. In this networks, message are rapidly shared by hubs and quickly distributed to subordinate nodes. This type of structure favors entrainment of columnar activity between local and distance columnar assemblies, that is, it favors the entrainment of the distinct *i*_*t, d*(*t*)_ s, what increases the correlation between 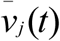 recorded by the different electrodes (Figure 4 and 5).

### 2.3 Cortical activity entrainment

The degree of cortical activity entrainment (**E**) depends on the characteristics of the activity in the columnar input fibers. For instance, thalamic oscillatory activity influences **E** using the Thalamic Reticular Fibers and it is responsible, among others, for the circadian **E** modulation; attention shifts, etc. Activity in sensory systems provides another source for controlling **E** and focusing cortical analysis of sensorial environment. Input activity from motor systems also modulates E to organize cortical processing of intended or running motor actions. In this line of reasoning, entrainment is the mechanism to enroll the adequate CCAs to solve complex tasks as proposed by the theory of distributed processing systems (e.g., Iturria-Medina et al, 2008; Ferrv et al 2008; Rocha, 1997 and Rocha et al, 2004 and 2011)

**E** is also influenced by columnar state governed by the distinct modulator neural systems: cholinergic, histaminergic, monoaminergic and serotoninergic systems. These circuits govern, among other, the degree and type of columnar cellular activity (e.g Rocha et al, 2001,2004 and 2011).

From discussion in the preceding section, the degree of **E** determines the degree of correlation (R) between the electrical activity recorded by the different electrodes. In this line reasoning, Rocha et al (Foz et al, 2002) proposed to use R to calculate the amount of information provided by each electrode about the entrained cortical processing. The rationality of this proposal is the following.

Let *R*(*i, j*) to be the calculate regression coefficient between 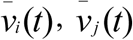 recorded by electrodes *e*_*i*_, *e*_*j*_ during a given EEG epoch selected to be studied. If these recorded activities are well correlated then *R*(*i, j*) tends to 1; if they are not correlated then *R*(*i, j*) is equal to 0. The maximum uncertainty about the relation between 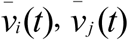 occurs when *R*(*i, j*) = 0.5. In the same line of reasoning if 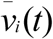 increases and 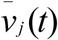 decreases (or vice versa) tend *R*(*i, j*) tends to −1. and the maximum uncertain occurs if *R*(*i, j*) = −0.5. Hence, given |*R*|(*i, j*) as the absolute value of *R*(*i, j*), the amount of uncertainty *H* (*i, j*) about the correlation between the electrodes *e*_*i*_, *e*_*j*_ increases as |*R*|(*i, j*) tends 0.5 and *H* (*i, j*) decreases if |*R*|(*i, j*) tends to 1 or 0. Because of this, Rocha et al (2001, 2017) proposed to calculated *H* (*i, j*) as

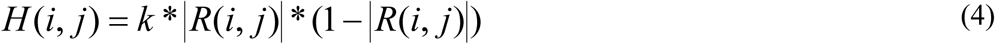

where *k* is a constant, such that if *k=*4 then maximum uncertainty is 1 when |*R*|(*i, j*) = 0.5.

By the same reasoning as above, if 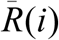 is the average of *R*(*i, j*) calculated for electrode *e*_*i*_ and all other remaining *e*_*j*_, then the amount of uncertainty 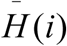 provided the electrode *e*_*i*_ about a cortical processing, tends to be minimum if 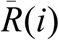 tend to 1 or 0 and maximum if 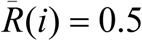. Thus

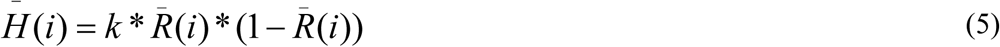

In this context, given the EEG recorded by a set of ***n*** electrodes, it is proposed that the amount of entrainment *E*(*i*) measured by the electrode *e*_*i*_ is calculated as the informational entropy

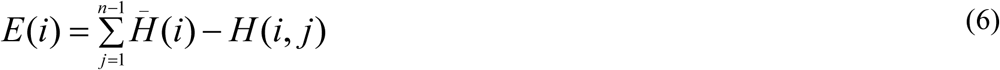

and the degree of entrainment *E*(*P*) of a recorded EEG epoch P is

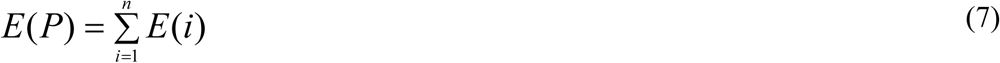

In the case of 20 electrodes, the maximum possible value of *E*(*i*) is 19 obtained when half of the correlation coefficient |*R*|(*i, j*) are 0 and the other half are 1. In this case 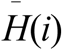 is 1 and *H* (*i, j*) = 0 for all other electrodes *e*_*j*_ because *R*(*i, j*) is either 1 or 0. In such a condition the maximum possible value for *E*(*P*) is 380.

The study of a cognitive task requires each volunteer of ***V*** volunteers (41 in Quiz task) to solve ***K*** (30 in Quiz task) activities composed of ***P*** phases (3 in Quiz task). In this condition it is necessary to calculate for each task phase ***P***:

1. the average *Ē*(*i*) of measured entrainments *E*(*i*) of each electrode *e*_*i*_ as

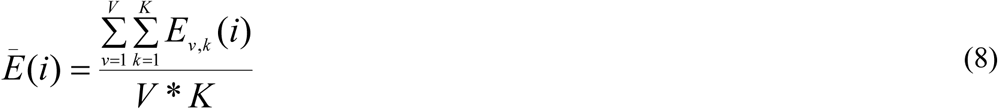

where *E*_*v,k*_ (*i*) is the entrainment of that electrode calculated for activity *k* and volunteer *v*; and
2. the mean *Ē*(*P*) entrainment calculated for phase ***P*** as

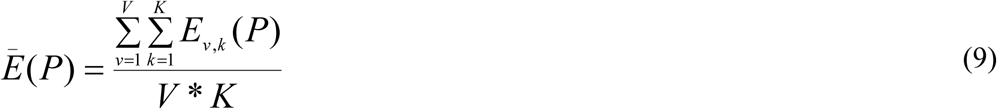

where *E*_*v, k*_ (*P*) is the entrainment calculated for activity *k* and volunteer *v.*

## 3 The EEG analysis

### 3.1 Entrainment brain mappings

Brain mappings in figure 6 were built color encoding the calculated *Ē*(*i*) for EEG epochs Q, S and C.. Maximum *Ē*(*i*) measurements are 2.62, 2.84 and 2.84 and minimum *Ē*(*i*) values are 1.69, 1.55 and 1.67 for Q, S and C, respectively. The corresponding *Ē*(*P*) values are 40.8, 40.5 and 42.5. These results shows that mean entrainment in the studied language processing is around 10% of the possible maximum entrainments. *Ē*(*i*) values are high for frontal electrode, manly FP1 and FP2, and they are low for posterior electrodes, manly PZ. *Ē*(*i*) values slightly differ between the epochs Q, S and C. Similar results were obtained in studies of other different reasoning processes (see Foz, 2002; Ribas et al, 2013; Rocha, et al 2010, 2001; 2013, 2014 and 2017; Vieito et al, 2013).

**Figure 6.**
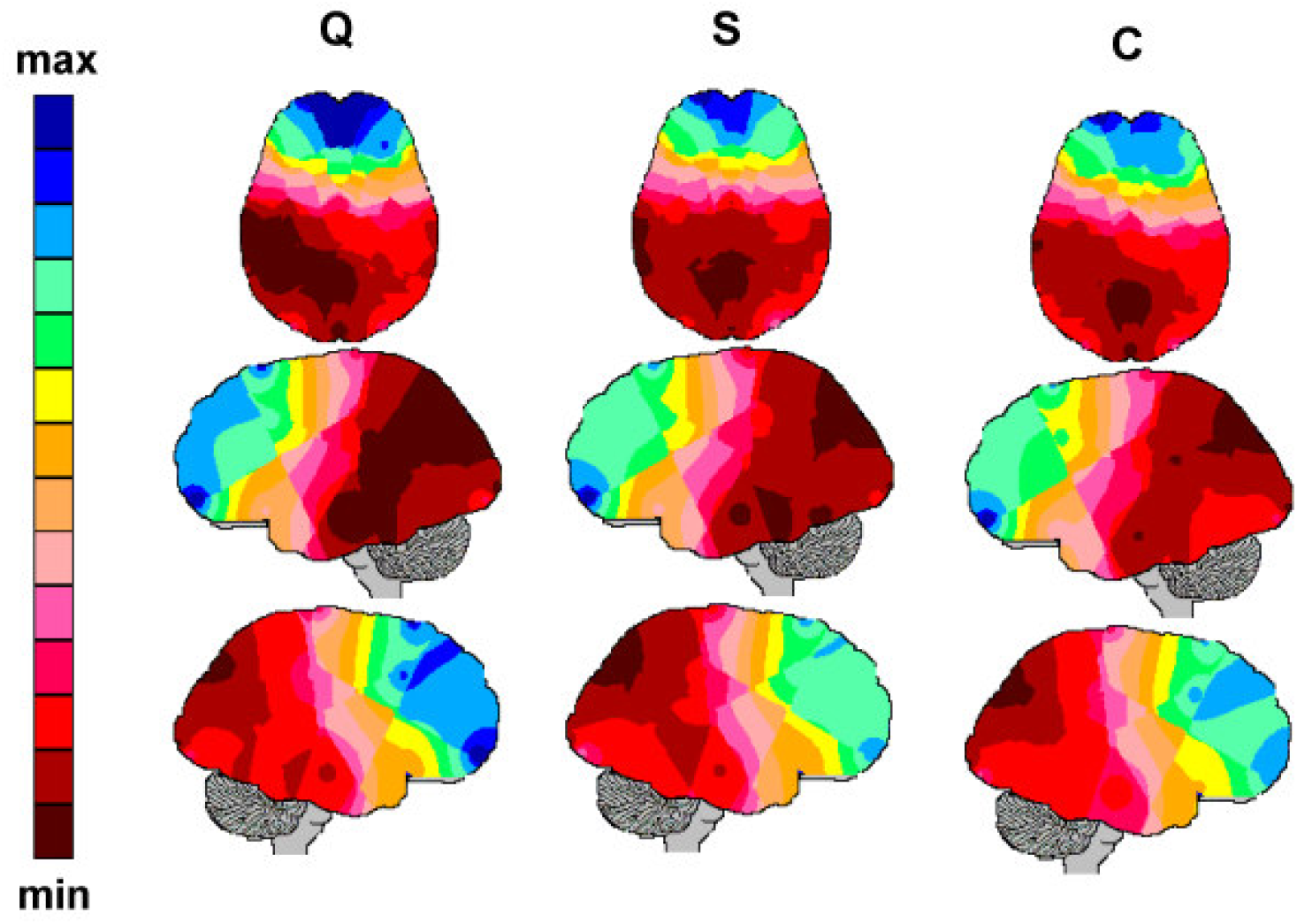
Entrainment mappings for EEG epochs Q, S and C. The values of *Ē*(*i*) were calculated from the regression coefficient *R*(*i, j*) calculated for each epoch (see Figure 5) for illustration of the regression coefficients calculated for epoch Q. *Ē*(*i*) values are normalized to built the mappings according to color code showed at left.

Because entrainment is favored in scale free networks due to the high connectivity of hub nodes, it is expected that *E*(*i*) distribution must follows a power law (Barabási and A. Réka, 1999. Iturria et al, 1998;). In other words, the frequency (*F* (*E*(*i*))) of each measured *E*(*i*) between 0 and max has to be high for low *E*(*i*) values and to rapidly (power law) decrease to very low for those *E*(*i*) close to max(*E*(*i*)), in other words

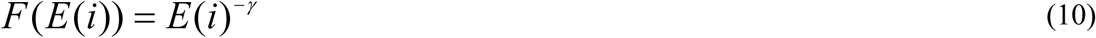

Figure 7 shows *F* (*E*(*i*)) vs *E*(*i*) relation in a log/log scale for EEG epochs Q, S and C. The fact that a liner function fit data with R^2^ greater than .7 and with γ around 2.4 confirms that *E*(*i*) distribution follow a power law (Iturria et al, 2008 and Rocha et al, 2011).

**Figure 7.**
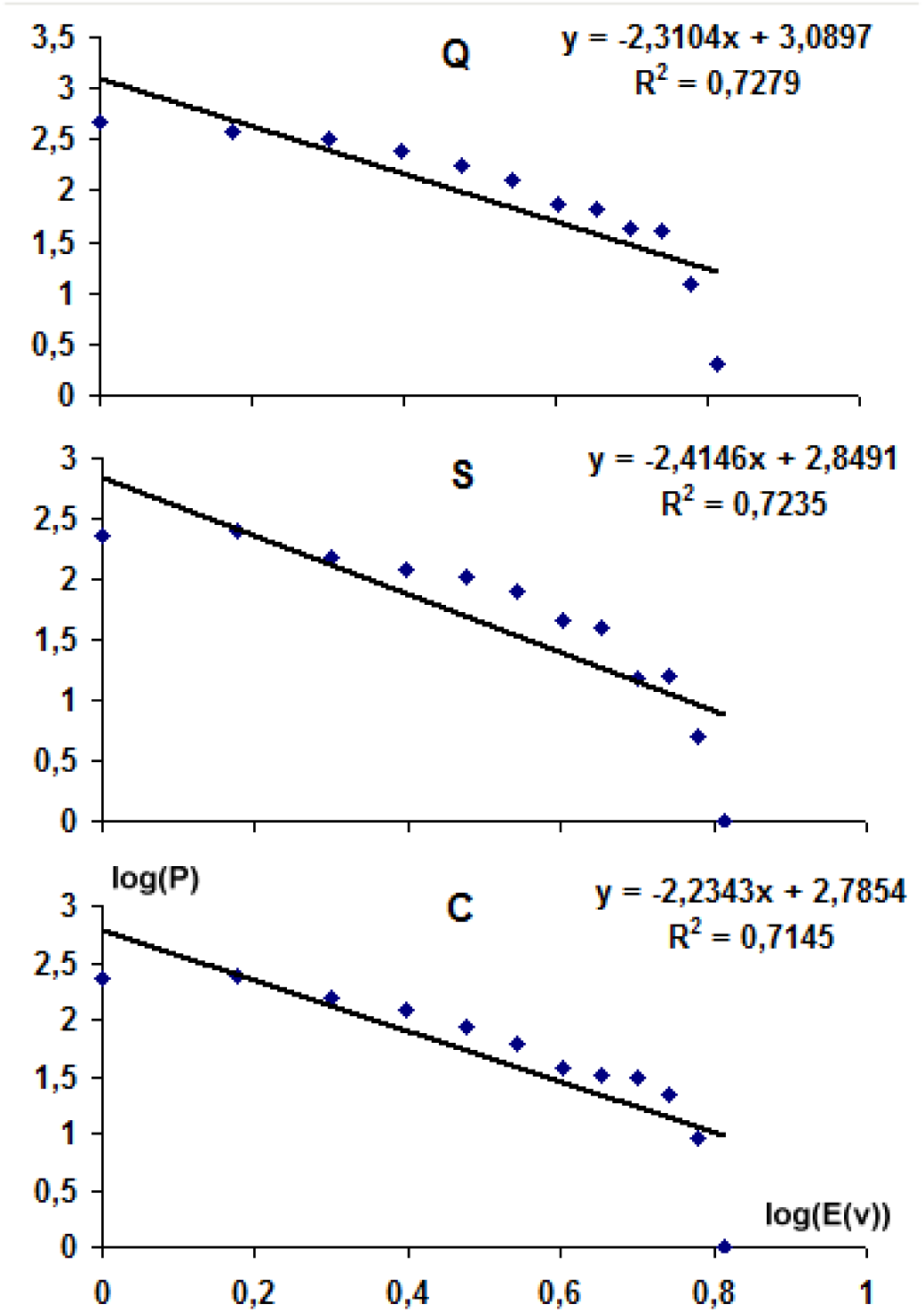
Entrainment *E*(*i*) scale free distribution. A scale-free network is a network whose degree distribution follows a power law, with 2< γ <3

### 3.2 Event Related Activity (ERA)

EEG_i_ data recorded during a selected EEG epoch (i = Q, S and C) of interest to study a given cognitive or reasoning task **T** (e.g., Quiz task) is composed by ***S*** samples of *v*_*n*_ (*s*) recorded at a specified rate (*R* hz) by each of the **N** recording electrodes. The value of ***S*** (512 in the Quiz task) is set by the sampling rate (e.g., 256 Hz in case of the Quiz task) and the duration of the EEG epoch (2s in the case of the Quiz task). Therefore, the EEG data base to be used to study **T** is composed by ***s*** lines of **N** measures *v*_*n*_ (*t*) for each EEG epoch selected to be studied. If each of the ***V*** (41 in Quiz task) volunteers solve ***K*** (30 in Quiz task) activities of **T** then the EEG data base will contain ***K*\**S*\**N*** measures *v*_*n, v, k*_ (*s*) for each selected EEG epoch. The following averages may be computed on this data base:

1. EEG_i_ average for task **T**, each electrode and each volunteer *v* is

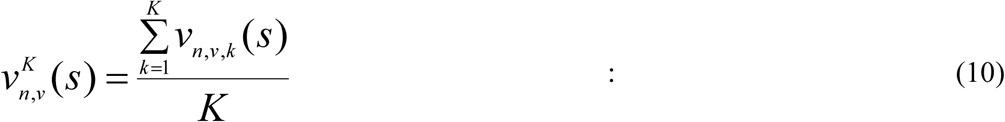
2. EEG_i_ average for task **T**, each electrode and taking into account all volunteers is

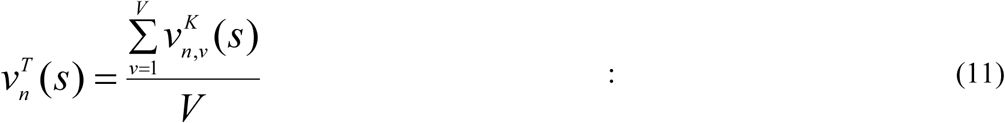
3. EEG_i_ average for task T and taking into account all recording electrodes is

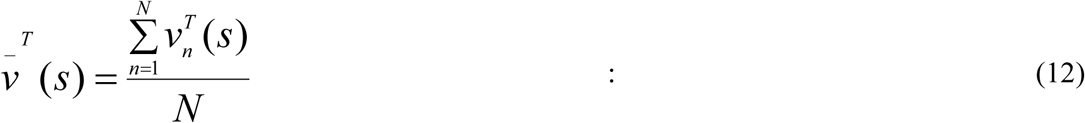

The different averages defined by equations 10 to 12 provides distinct pieces of information about the cortical electrical activity of a Event Related Activity 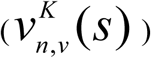 defined for each selected task EEG_i_ epoch: electrode and volunteer (eq. 10); 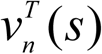 defined for each selected task EEG_i_ epoch, and electrode, and 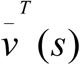 defined for each select task EEG_i_ (eq. q12) epoch. Figure 8 display the evolution of 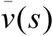 for each EEG epochs in case of the Quiz task.

**Figure 8.**
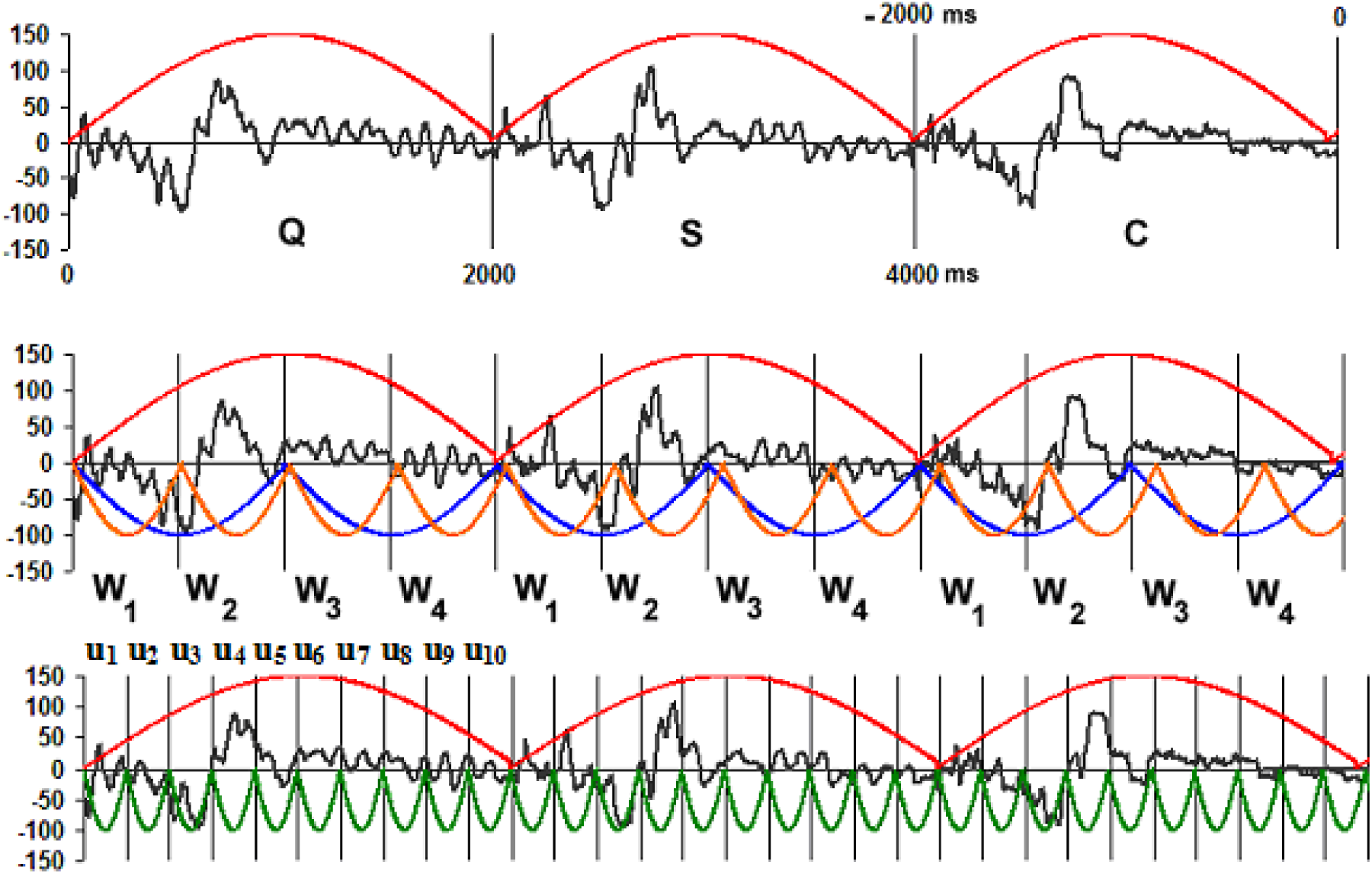
Grand average calculated for EEG epochs Q, S and C. Four distinct waves is evidenced by segmenting ERA_i_ with the 2 Hz sinusoidal function in blue. In addition, the 1 Hz ERA_i_ segmentation isolates two distinct components. the first one (W_1/2_) enveloping W_1_ and W_2_ components in Figure 2 and the second one (W_3/4_) enveloping W_3_ and W_4_ in Finally, the 5 Hz ERA_i_ segmentation isolates clear 5 subcomponents (v_j_) of W_1/2_ and possible 5 subcomponents (v_j_) of W_3/4_.

#### 3.2.1 ERA waves

Many distinct ERA components have being identified in the reductionist approach, by its polarity and time occurrence: eg: N50 - negative wave peaking at 50 ms after stimulus onset; P100 - positive wave peaking with a latency of 300 ms; N400 - negative wave peaking at 400 ms; P600 - positive wave peaking at 600 ms, etc. (e.g., Donchin, et al 1975; Kutas, and Hillyard, 1980; Morgan; Hansen and Hillyard, 1996 and Picton, et al 1974 *and* Regan,1966). However, if the recorded *v*_*n*_ (*s*) is generated by a complex time locked set of oscillatory activities produced by many different CCA sets, then, instead of searching for specific *v*_*n*_ (*s*) at defined samples ***s***, it is necessary to identify possible characteristics ERA waves that may result from a concerted synchronized activity.

Calculated Quiz ERA_i_ (i= Q, S and C) are shown in Figure 8. Its inspection revealed that these averages are nicely delimited by the .5 Hz sinusoidal function (pink). In addition, these ERA_i_ are constituted by four distinct EEG waves (W_1_ to W_4_) of approximately 500ms of duration as revealed by the 2Hz sinusoidal segmentation function (red). W_1_ is a negative wave occurring at the beginning of each ERA_i_. W_2_ is a predominant positive wave following W_1_. W_3_ is composed by a series of positive peaks occurring from 1000 to 1500ms after the beginning of each ERA_i_. Finally, W_4_ is composed by a series of positive and negative peaks following W_3_.

Two other important ERA_i_ segmentations are provided by the 1 (blue) and 5 (green) Hz segmentation functions in Figure 8. The components isolated by 1 Hz function are totally different: the first one (called here W_1/2_ component) combines components W_1_ and W_2_ isolated by the 2Hz function and the second one (called here W_3/4_ component) is composed by the components W_3_ and W_4_ isolated by the same 2Hz function. The components (called here u_j_ isolated by the 5Hz function are more variable within each ERA_i_ and between the different ERA_i_. They are mostly composed by small biphasic oscillations superimposed upon the 2 Hz components W_i_.

Volunteers took around 2 seconds (Rocha et al, 2015) to make picture choice what may explain the similarity between the 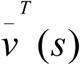 calculated for tasks S and C, if it is assumed that picture selection occurs just after the end of visual processing during S. This visual processing started, after Quiz listening or reading, with visual picture display and ended with picture selection, therefore it is correlated to the recorded activity during EEG epochs S and C. If this visual processing time is assumed to take around 2 seconds for all volunteers, then choice picture selection is synchronized with picture display with a delay of 2000 ms. Regression analysis showed that ERA_i_ in Figure 8 are correlated with a R around 0,8, reflecting the dependences of the reasoning discussed above.

Oral quiz texts were recorded by a native Brazilian speaker instructed to produce a clear speech to avoid any misunderstanding of the quiz that could jeopardize the experiment. No volunteer complained about the quiz understanding. Measurement of the duration of these recorded texts revealed a mean duration around 2000 ms. Because of this, the display of the written texts were fixed in 2500ms. Taking these facts into account, Rocha et al (2015) and Pereira et al. (2017) interpreted the similarity of 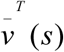 calculated for EEG epochs Q and S as due to a modular cortical processing of 2000 ms composed by 4 main steps disclosed by the identified W_i_s. Similar modular cortical processing was observed for other types of reasoning studied in our group (Rocha et al, 2017), too

#### 3.2.2 Identifying ERA waves

A polynomial is a an expression consisting of variables and coefficients that involves only the operations of addition, subtraction, multiplication, and non-negative integer exponential variables (Childs, 2009). More precisely, it has the form:

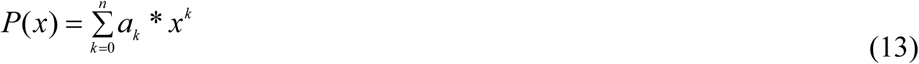

where *coefficients a*_*k*_ may be integers, rational numbers, real numbers, complex numbers or, more generally, members of any field.

The number of exponential variables determines the polynomial degree (*n*^*th*^ degree in the above equation). If the leading coefficient *a*^*n*^ is positive then the polynomial function increases to positive infinity at both sides and thus the function has a global minimum. Likewise, if *a*^*n*^ is negative, the polynomial decreases to negative infinity and has a global maximum.

The complexity of a polynomial is determined by the number: 1) of their critical points at which the polynomial derivative is zero; 2) global maximums and minimums and 3) real roots or zero crossings.

Many n^th^ Polynomial regression (nPPR) may calculated for each ERA_i_ to identify its possible different waves (W or u). This regression is calculated by computing a regressed *v*_*r*_(*t*) as

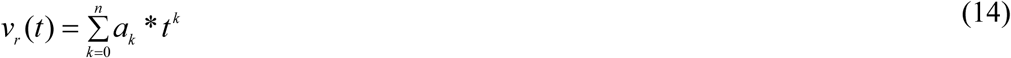

and calculating its correlation to the averaged *v*(*t*). Although polynomial regression is technically a special case of multiple linear regression, the interpretation of a fitted polynomial regression model requires a somewhat different perspective. It is often difficult to interpret the individual coefficients in a polynomial regression fit, since the underlying monomials can be highly correlated. It is generally more informative to consider the fitted regression function as a whole.

Figure 9 shows three (n=20, 12 and 6) different nPPRs calculated for Q, S and C ERAs used to identify a possible common ERA pattern. The best fitting is provided by the 20 PPR, that reveals that the all global ERA_i_ waves are composed by 13 critical points (where the derivative is zero). 6 global maximums, 7 minimums and around 6 real roots (0 crossings).

**Figure 9.**
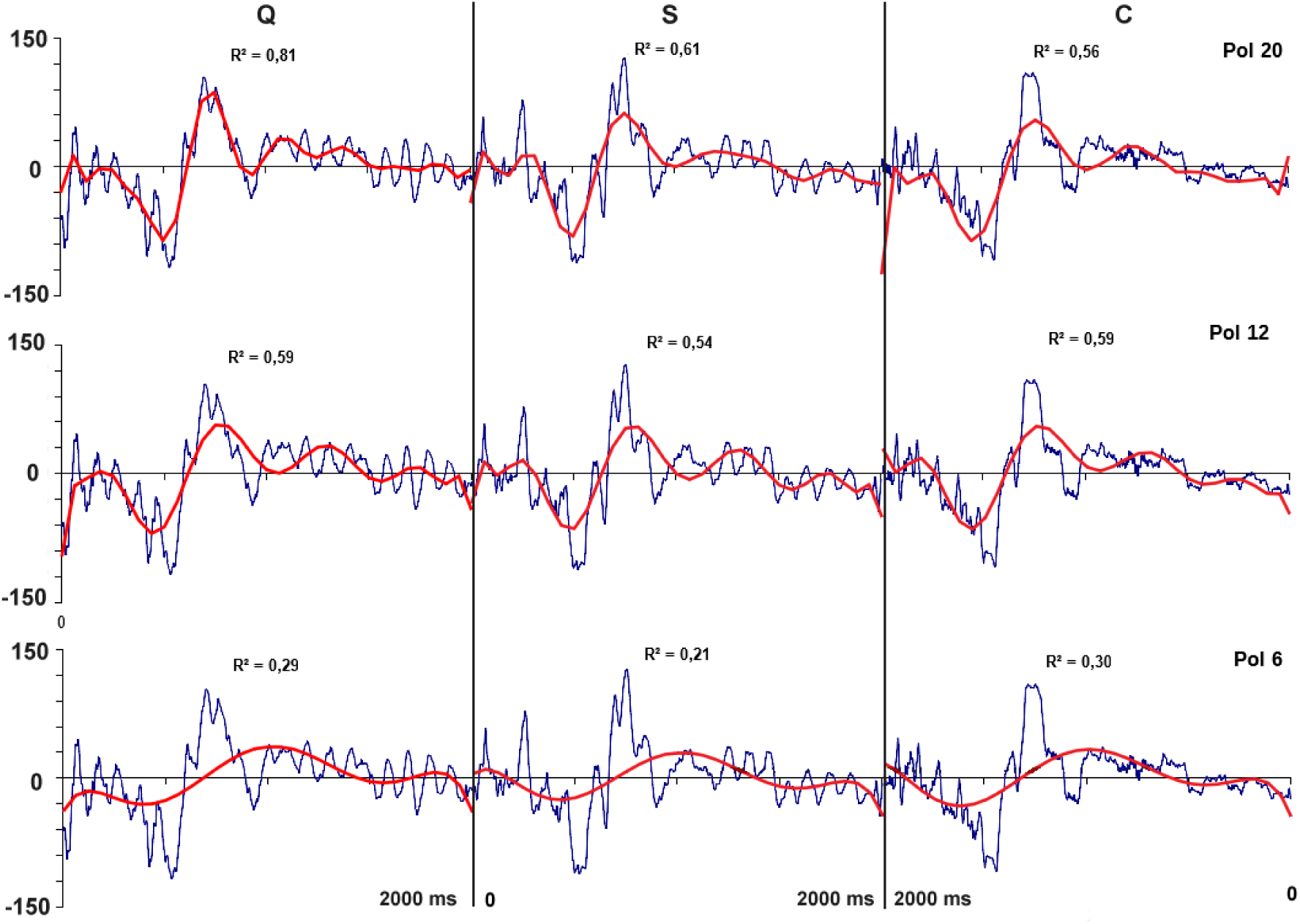
Polynomial regression (red) calculated for different (20, 12 and 6) polynomial degrees for the different Q, S and C ERAs (blue). The regression square coefficient R^2^ decreases with the polynomial degree for all ERAs, being highest for n=20. R^2^ varies from ERA Q to C, but disclose a general common pattern for all these ERAs. In case of 20 PPR, this pattern involves around 13 critical points (where the derivative is zero). 6 global maximums, 7 minimums and around 6 real roots (0 crossings).

All the different ERA_i_ as modeled by similar polynomials, independent of their degrees (20, 12 and 6). This suggested that ERA_i_ are similar for the EEG_i_ epochs Q, S and C.

Figures 10 and 11 shows the nPPR modeling as above for ERA components W_1/2_ and W_2/3_ disclosed by the 1 Hz sinusoidal segmentation function (Figure 10) and components W_i_ disclosed by the 2 Hz sinusoidal segmentation function. The results of these nPPR modeling add more information about how to use this type of fitting to identify ERA waves.

**Figure 10.**
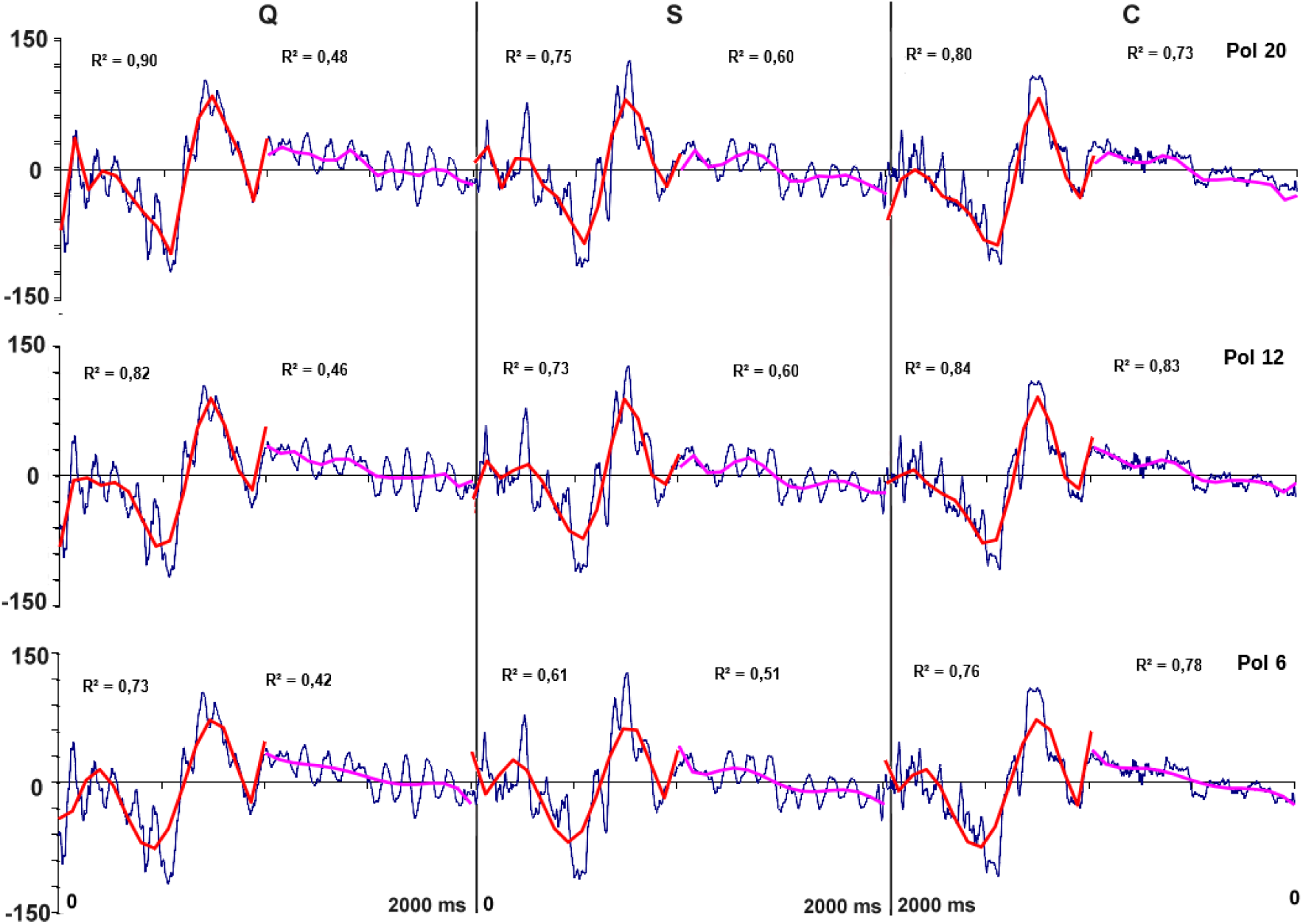
Polynomial regression (red) calculated for different (20, 12 and 6) polynomial degrees for the different Q, S and C ERAs components W_i/i+1_ (blue). The regression square coefficient R^2^ decreases with the polynomial degree for all ERAs, being highest for n=20. However, these coefficients are very close to each other when 20 and 12 PPR are compared.

**Figure 11.**
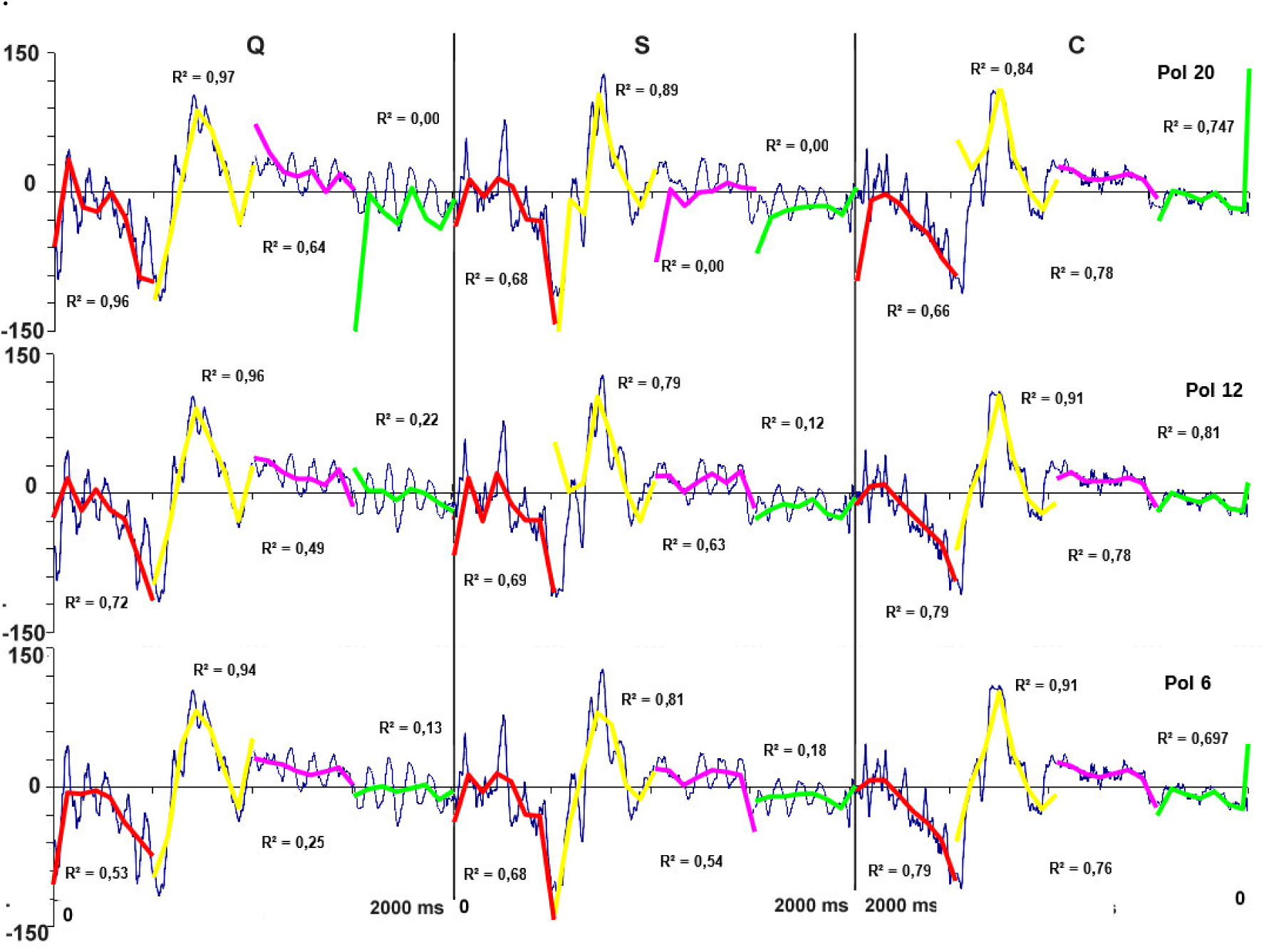
Polynomial regression (red) calculated for different (20, 12 and 6) polynomial degrees for the different Q, S and C ERAs components W_i_ (blue). The regression square coefficient R^2^ are almost invariant with the polynomial degree for all ERAs.

Inspection of Figure 10 shows that both 20 and 10 PPR calculations result in very similar R^2^ values, indicating that increasing polynomial complexity in this case does not add much more information about ERA waves that may justify the cognitive effort of reasoning with more complex polynomials. The difference in explaining more data variance using 10 PPR instead of 6 PPR may, perhaps, justify the choice of the 10 PPR modeling to identify components W_1/2_ and W_3/4_. But this choice depends on how precise someone wants to be in discussing ERA components.

Results displayed in Figure 11 about W_i_ nPPR fitting, clearly shows that, in some cases, increasing complexity of nPPR modeling, does not add any additional relevance piece of information. This is because variation of R^2^ obtained for the different used polynomial degrees, did not change significantly the amount of explained data variation. In this case, it is advisable to keep analysis as simple as possible, opting for the simplest nPPR modeling or the 6 PPR fitting.

#### 3.2.2 Isolating and identifying other ERA waves

Once the 6 PPR nicely fits the ERA_i_ components W_i_, it is possible remove these components from the calculated ERA, in the attempt to better visualize other smaller waves. This is done by subtracting the regressed *v*_*r*_ (*t*) from the averaged *v*(*t*). The graphic ERA & PPR in Figure 12 shows, for all EEG epochs, the computed *v*(*t*) (blue) and *v*_*r*_ (*t*) (different colors) calculated for 6 PPR, whereas the graphic ERA - PPR display the corresponding differences *v*(*t*) − *v*_*r*_ (*t*). The 5 Hz segmentation function (green in ERA-PPR graphic) discloses a set of small waves that are nicely fitted by the 6 PPR modelling (red in Q ERA - PPR & Pol 6 graphic). The results of this fitting for Q ERA_i_ are shown in Figure 12 and identifies the waves labeled u_1_ to u_10_ with a degree of accuracy (R^2^) greater than .8. Results for the other EEG epochs S and C are similar to those displayed for epoch Q. The characteristic pattern of these waves is composed by a 10 Hz activity as shown in *u pattern* in Figure 12.

**Figure 12.**
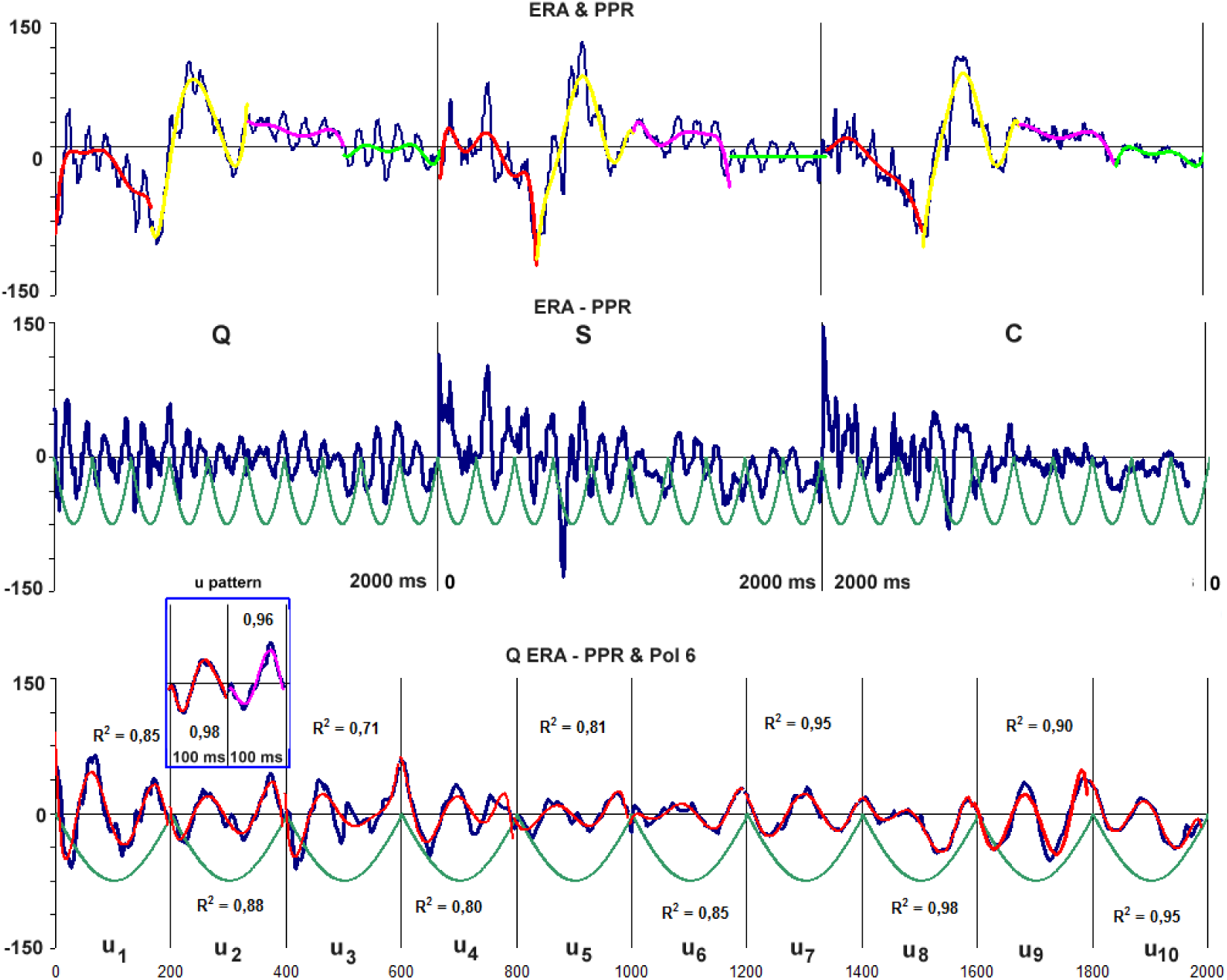
ERA theta segmentation and u_i_ components. Graphic ERA & PPR display the regressed *v*_*r*_ (*t*) (various colors) from the averaged *v*(*t*)(blue) for EEG epochs Q, S and C. Graphic ERA - PPR displays the corresponding differences *v*(*t*) − *v*_*r*_ (*t*) (blue) and the waves u_i_ disclosed by the 5 Hz segmentation function. Graphic Q ERA - PPR & Pol 6 displays, for EEG epoch Q, the calculated *v*(*t*) − *v*_*r*_ (*t*) (blue) and their corresponding regressed values (red) for 6 PPR. The 6 PPR modeling fits data with R^2^ greater than .8. A 6 PPR fitting for a 10 Hz segmentation is inserted in the figure labeled as *u pattern* to illustrate the general characteristic of the u_j_ waves.

PPR results displayed in Figures 9, 10, 11 and 12 shows that polynomial regression is very useful tool for nicely and formally characterizing all ERA_i_ waves visually identified in Figure 8 with the aid of all sinusoidal segmentation functions.

### 3.3 ERA oscillatory activity

As pointed before, different oscillatory circuits contribute to the genesis of the cortical electrical field recorded by the distinct electrodes of a EEG recording system. Fast Fourier Transform (FFT) is used to decompose the recorded EEG into the set of sinusoidal functions associated to these oscillatory cortical activity. This type of analysis produces the frequency cross spectrum (frequently labeled CRS) showing the power each sinus wave contributes to the genesis of the recorded EEG.

Figure 13 shows the CRS calculated for the EEG epochs Q, S and C in the range of 1 to 64 Hz. These CRS are distinct for each of the studied epochs, showing a predominance of low frequencies, specially the 5 Hz one in case of epoch Q; a predominance of high frequencies, specially the 64 Hz one in the case of epoch S, and the great predominance of 5 Hz frequency in case of epoch C. CRS of this last epoch is very similar to that calculated for epoch Q and distinct of that obtained for epoch S.

**Figure 13.**
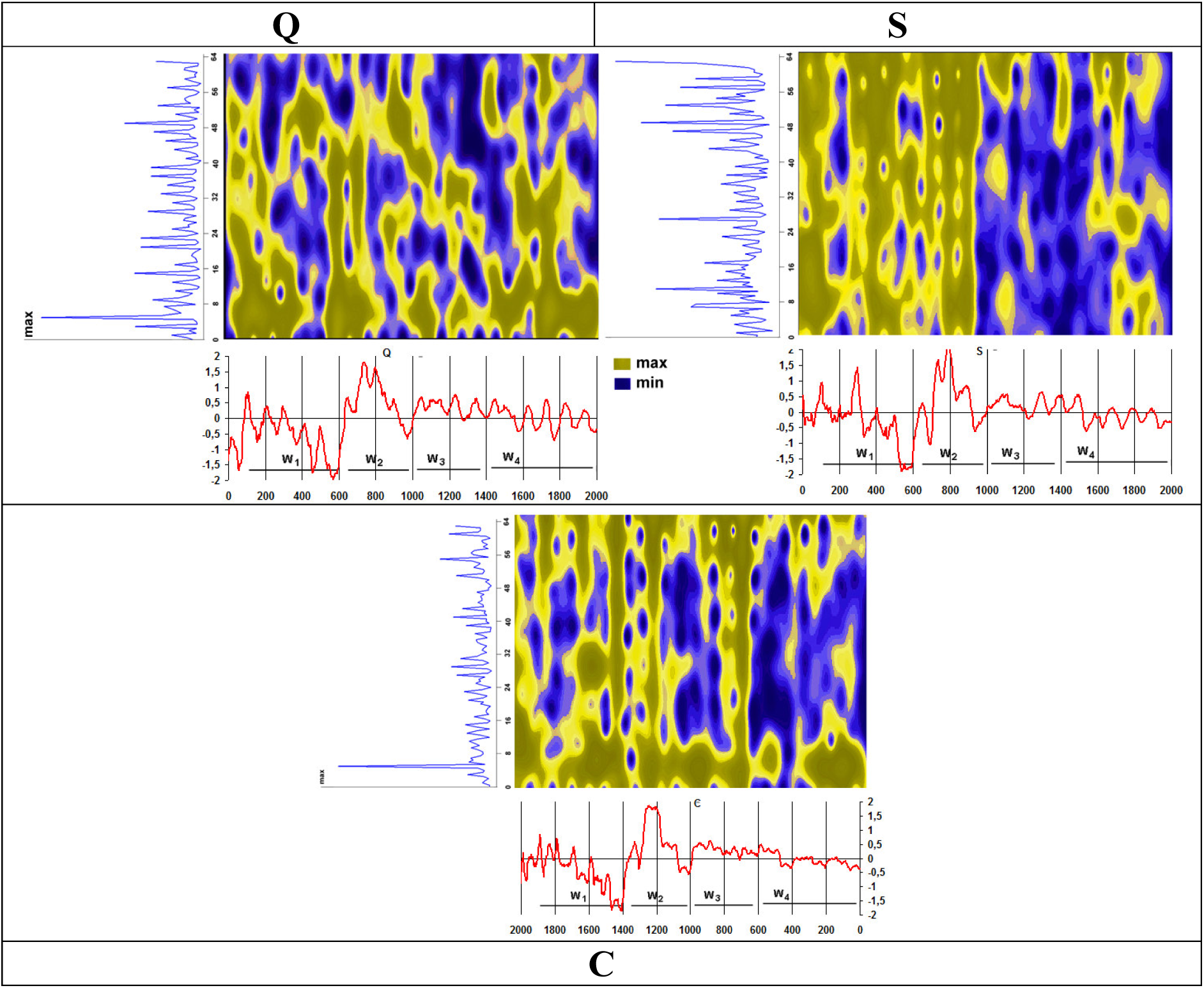
ERA oscillatory activity for EEG epochs Q, S and C. Each graphic show the cross spectrum (CRS) at upper left, time varying cross spectra (TVCRS) at upper right and Grand average at bottom. Cross spectrum results shows that many different oscillatory activities are important contributors to the recorded EEG. In addition, TVCRS results show that many different oscillatory activities contribute to generated the each sampled *v*_*n*_ (*s*)

It is possible to study time evolution of the contribution of each oscillatory activity to the ERA genesis. One of the techniques available is the Time Varying Cross Spectrum (TVCRS) and EEG wavelet analysis is another popular technique. Here, the first one was used as implement in the LORETA software freely available in Internet^1^. This technique calculates TVCRS for all sampled 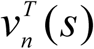 using a specified window (here, 50 ms). For further discussion about TVCRS see Rocha et al (2017).

Figure 3 shows that TVCRS calculated for EEG epochs Q, S and C are very different. Despite this, some general patterns arise from inspection of the different TVCRSs. The first one is the contribution of the majority of the studied frequency to genesis of W_1_ and W_2_ no matter the EEG epoch. Another general feature is the smaller number of the frequencies contributing to W_3_ and W_4_. Low frequencies (smaller than 12 Hz) are continuously active during the duration of epochs Q and C, while high frequencies (greater than 50 Hz are activated during the entire epochs S and C. However, the most interesting result from TVCRS analysis is the fact that many distinct oscillatory activity concurrently contribute to each recorded sample *v*_*n*_ (*s*) as proposed, e.g,, Burgess (2012) and Klimesch et al (2007).

Although ERA_i_ calculated for EEG epochs Q, S and C are visually very similar and highly correlated, spectral analysis of their associated oscillatory activity clear differentiates each ERA_i_. This stress the necessity of using all available techniques if one wants to have a realistic understanding of the recorded EEG and the underlying cortical activity.

### 3.4. Factorial Analysis of cortical entrainment

Factor Analysis (FA) is a statistical tool to investigate patterns *P*_1_ of covariation in a large number of variables and to determine if information may be condensed into small sets of these variables called principal components *P*_*i*_ (Hair et al, 1998 and Rocha et al, 2017). This transformation is defined in such a way that the first principal component *P*_1_ is the one that accounts for as much of the variability in the data as possible, and each succeeding component (*P*_2_, *P*_1_, …) in turn explain the subsequent amount of variance possible under the constraint of being orthogonal to (i.e., uncorrelated with) the preceding components.

FA may be used to study entrainment (*E*(*i*)) covariation during a cognitive task solution (Rocha et al., 2017). In case of the Quiz task, four patterns *P*_*j*_ explains 80% of *E*(*i*) covariation during all studied EEG_i_, *E*_*v,i*_ (*i*) calculated for each electrode *e*_*i*_, volunteer **v** and EEG_i_ has a specific contribution to each *P*_*j*_, called loading factor *f* _*j*_ (*e*_*i*_) that ranges from 0 (no contribution) to 1(full contribution). FA mappings of the cortical entrainment quantified by *E*(*i*) are computed taking into consideration these loading factors *f* _*j*_ (*e*_*i*_) that are color encoded to associated each *P*_*j*_ to the cortical area near the electrodes *e*_*i*_ heavily loading on it. Figure 14 displays these mappings calculated for EEG epochs Q, S and C and shows that they are very similar for all studied EEG epochs.

**Figure 14.**
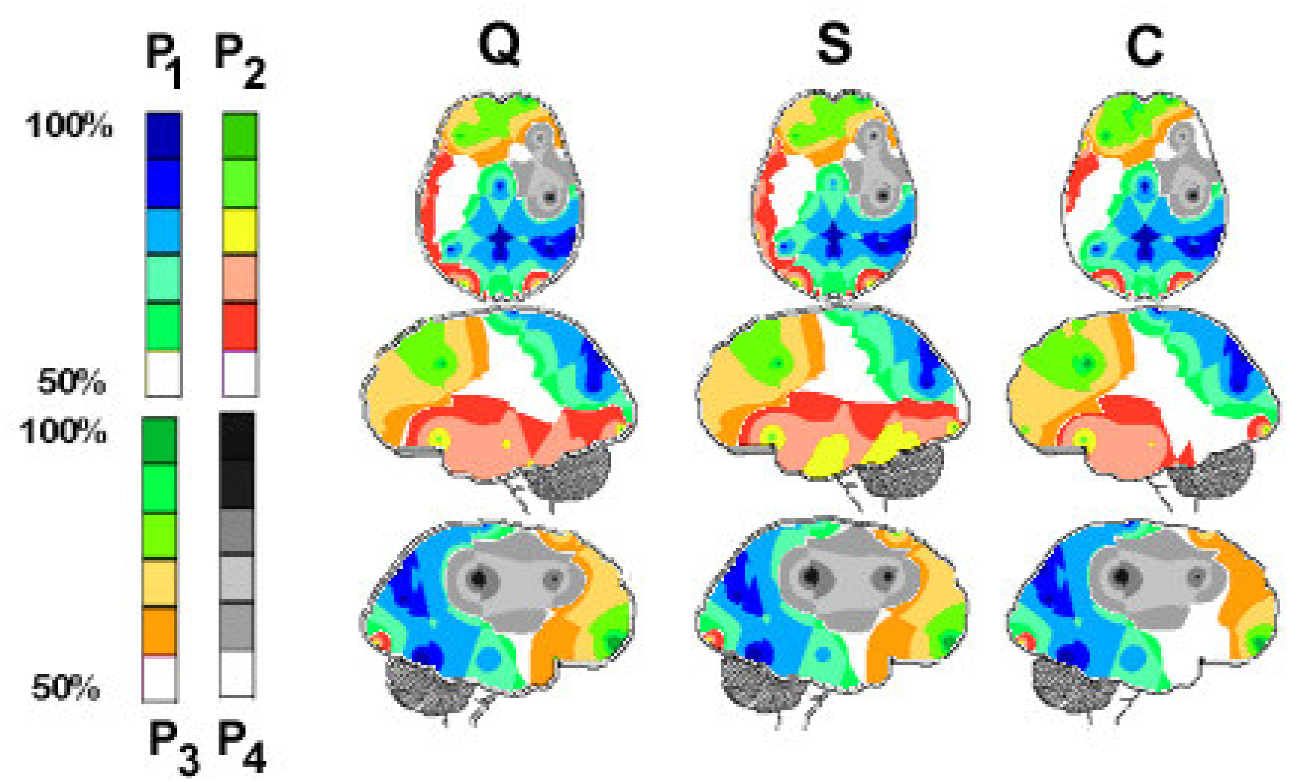
Factor mappings calculated for EEG epochs Q, S and C. Four different patterns *P*_*j*_ explained 80% of *E*(*i*) covariation. The loading factors *f* _*j*_ (*e*_*i*_) on these patterns *P*_*j*_ were used to produce the brain mappings according to the color codes at left.

Pattern *P*_1_ is composed by the entrainment of the activity recorded by the electrodes CZ, P3, PZ, P4, T4 and T6. Pattern *P*_2_ shows that entrainment of the activity recorded by the electrodes F7, T3, T5, O1 and O2 covaried together during epochs Q and S, whereas electrode T5 did not contributed to this pattern during epoch C. Entrainment of the activity recorded by the electrodes FP1, FP2, F3, FZ and F8 covaried together as disclosed by pattern *P*_3_. Finally, pattern *P*_4_ shows that entrainment of activity registered by electrodes F4 and C4 are correlated. Rocha et al (2015) interpreted these results assuming that pattern *P*_1_ disclose the entrainment involved in correlating verbal and visual information in order to solve the quiz. Pattern *P*_2_ was proposed to reflect entrainment of verbal cortical areas in decoding the quiz texts. Pattern *P*_3_ was considered to be associated to entrainment of frontal areas involved in controlling reasoning to solve the quiz. Finally, pattern *P*_4_ would disclosed entrainment involved in selecting the correct picture.

### 3.4 Identifying the columnar assemblies generating the recorded EEG

LORETAs (Pascual-Marqui et al., 2002a, b) was used to solve the inverse problem of identifying the cortical columnar assemblies activated during EEG epochs Q, S and C. Although LORETAs allows the selection of up to 5 of the most probable source generating the recorded *v*_*n*_ (*s*), it provides accurate x,y,z coordinates only for the first inverse problem solution. Because of this, we took for further analysis just the results of this first solution for the 512 *v*_*n*_ (*s*) samples during EEG epochs Q, S and C. Spatial location of these EEG Loreta Identified Sources (ILS) are shown in the brain mappings and their frequencies are displayed in the corresponding histograms of figure 11. Statistical significance threshold for Z score calculated for *v*(*s*) in case of 20 recording electrodes (as in present case) is 1.91. Because of these, only dos ILSs calculated for statistically significant 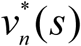 (Z score > 1.91) are selected for further analysis.

For all studied EEG epochs, ILS are widely distributed over the cortex, although they predominate over the frontal and occipital lobes. Visual inspection of data presented in figure 15, shows that ILS distribution differed for each studied EEG epochs mostly in case of those sources located at the parietal and temporal lobes. The most frequent ILS are located at BA 18 and 19 and their frequencies differ for each studied epoch. Sources located at BA 10 and 11 were also very frequent during epoch Q but reduced during epochs S and C. Sources located at BA 47 Inferior Frontal Gyrus enhanced their activity from epoch Q to S and from S to C. Here, it will be assumed that these ILSs are those generated by the columnar input activity arriving at the identified cortical area and generating *i*_*i*_ (*t*) (see section 2).

**Figure 15.**
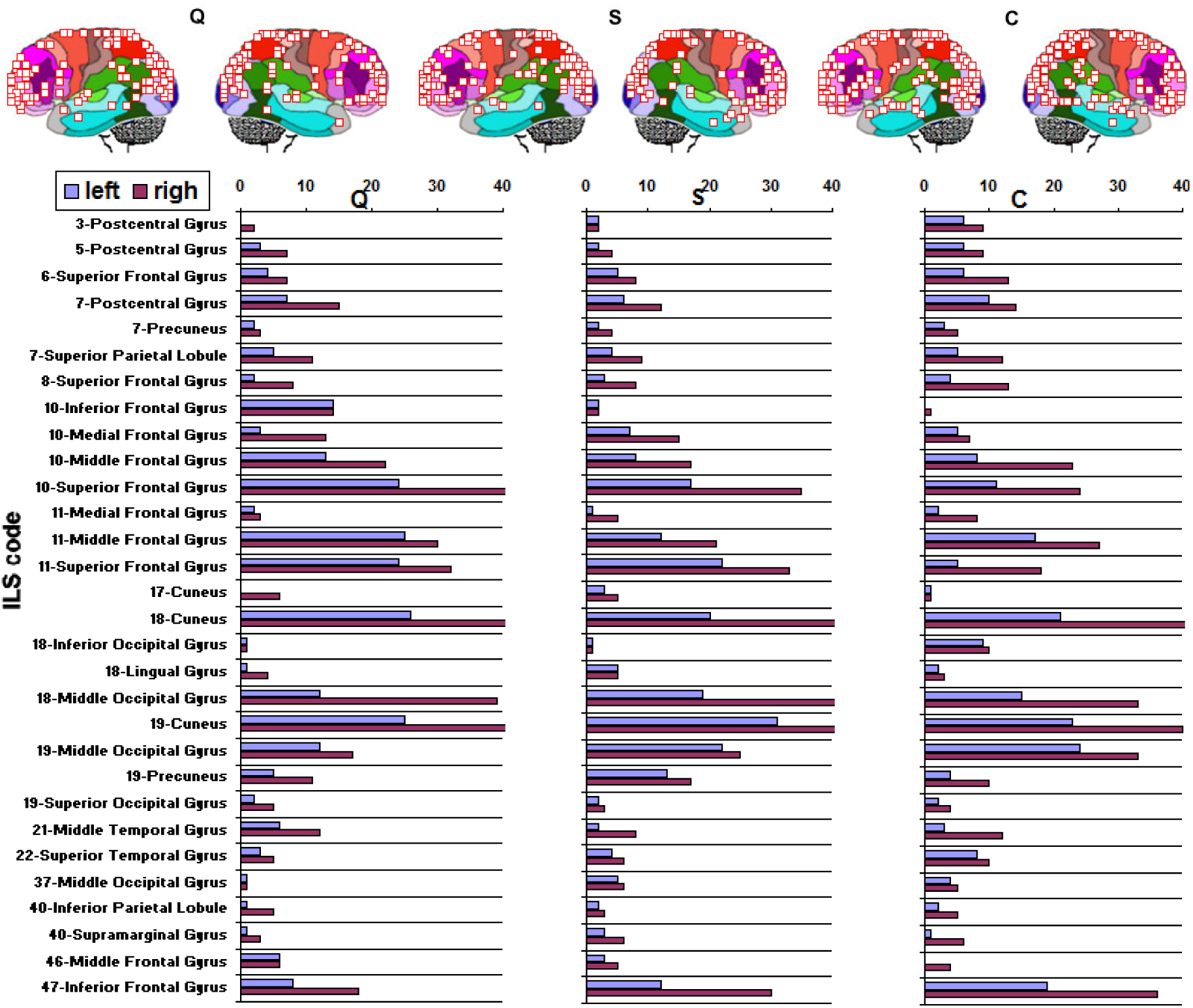
Spatial distribution (SLF) of significant LORETA identified sources (ILS) during epochs Q, S and C. ILS are coded taking into consideration their Brodmann area number and anatomic gyrus name. This code is named here ILS code. Results shows that many sources are common generators for all Q, S and C epochs being responsible for the similarity of the corresponding ERA_i_. The frequency each of these sources contributes to the recorded EEG varies for Q, S and C epochs, partially explaining the ERA_i_ observed differences.

Identified CCAs may be encoded by their BA area number and anatomic gyri (ILS code) as show in Figure 15. We identified, up to the moment, 114 locations in all decision making studies made by our group (see e.g. Rocha et al, 2017). This SLF encoding turns distribution of the identified CCAs a spatial ordered series and it permits the use of correlation to compare the different SLF_Q_, SFL_S_ and SFL_C_ distributions shown in figure 12. This analysis revealed that SLF_Q_ for EEG epoch Q correlated to SFL_S_ for EEG epoch S with a degree R=0.83; SLF_Q_ correlates to SFL_C_ with R=0.59, and SFL_S_ SLF(S) correlates to SFL_C_ with R=0.74. This provides a clear measure of the differences pointed above from visual inspection of figure 15.

Hub nodes are heavily connected to other nodes of a scale free and scale free connections rapidly decreases if nodes are adequately ranked. Entrainment *E*(*i*) was shown to be supported by a scale free network in section 4. In this condition, SFL is expected to be influenced by network connectivity and to increase with *h*(*e*_*i*_). This ranking provides a measure of node *hubness*. SLFs for EEG epochs Q, S and C were ordered in a increasing fashion and the corresponding order index was assumed to be a relative measure of hubness. Regression analysis disclosed a exponential relation between SLF and *h*(*e*_*i*_) as shown in figure 16. Hubness is high for CCA located at BA 10 or 11, BA 18 or 19 and BA 47 no matter the considered EEG epoch (compare Figures 15 and 16).

**Figure 16.**
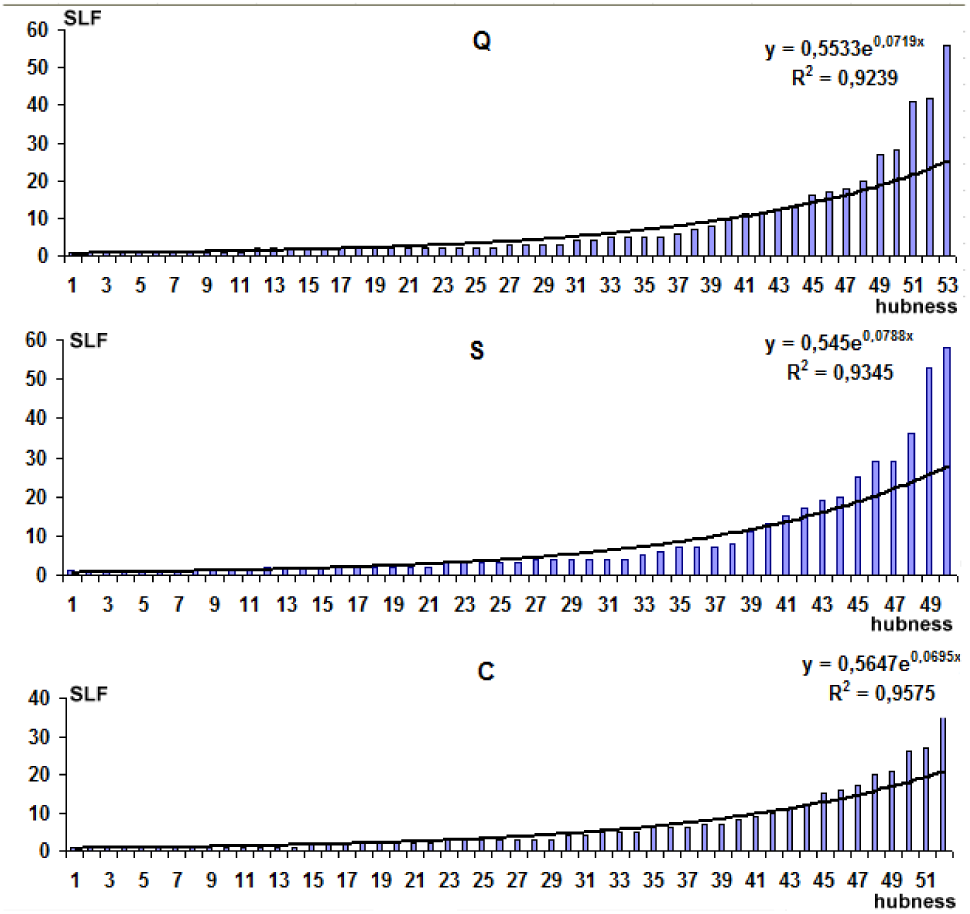
Hubness - SLF distribution follows a exponential law when is ordered in a increasing fashion and plotted according to the corresponding order index or hubness. Uses SFL value in Figures 11 and 12 to correlate ILS in Figure 2 to its hubness. For instance, Cuneus at BA 19 has hubness index of 53 in Q, 50 in S and 50 in C.

Figure 17 displays the ILS temporal distribution ITD(W_i_) associated to each wave component W_i_ and each studied epoch and Table 1 shows the calculated R for comparison ITD(W_i_) calculated for SLF_Q_, SLF_C_ and SLF_S_. These results show that SLF greatly varied for each W_i_ component taking into consideration the distinct studied EEG epochs.. In addition, Table 2 shows the calculated R between the different ITD(W_i_) showing that SFL greatly varies for the different components W_i_.

**Figure 17.**
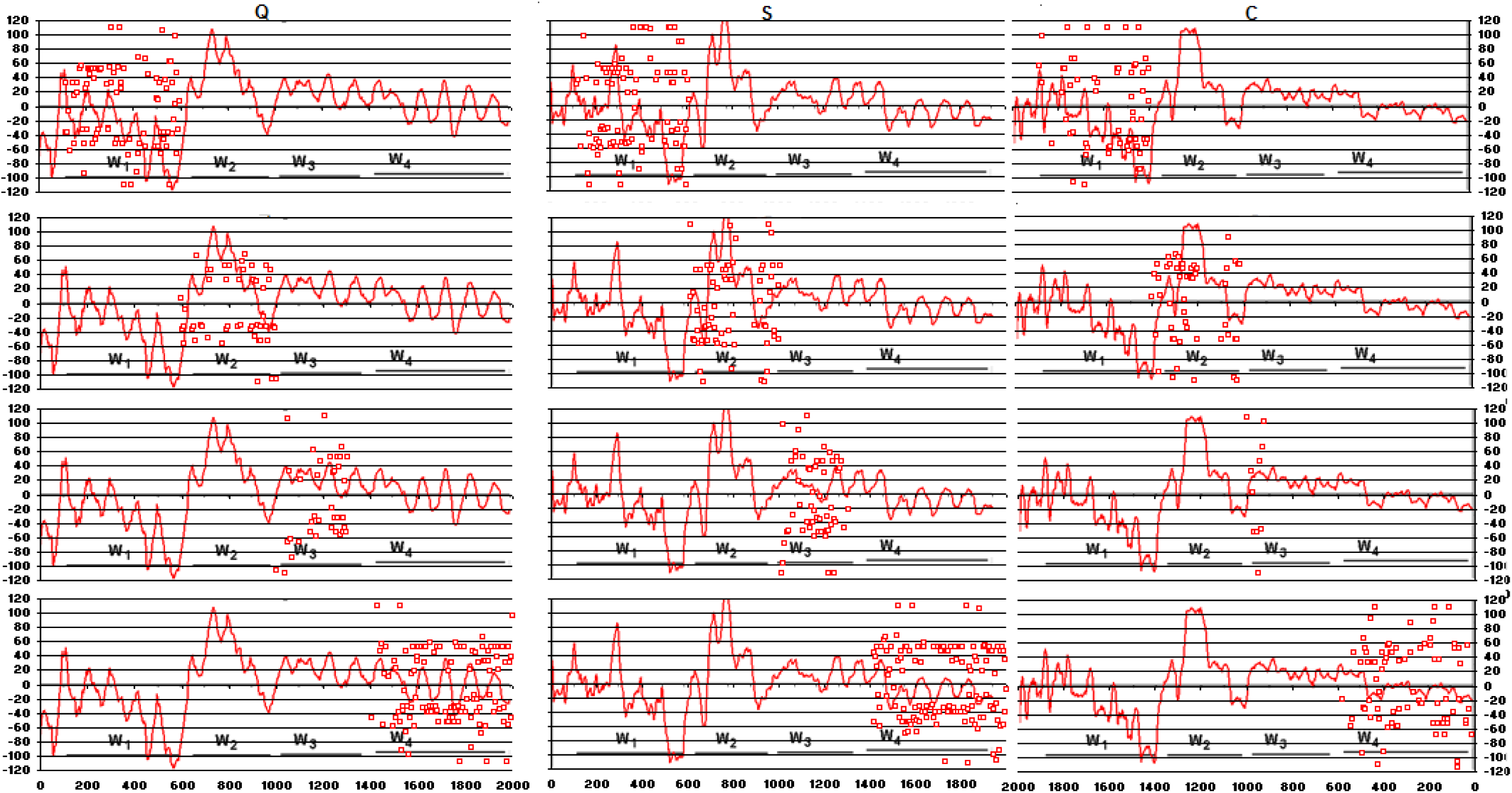
Temporal ILS and EEG components W_i_ for epochs Q, S and C

The following conclusions derive from data in figure 11, 12 and 13 and tables 1 and 2:

**Table 1.** R values calculated for SLF epoch relations and each W_1_.

| | $SLF_Q/SLF_S$ | $SLF_Q/SLF(C)$ | $SLF_C/SLF_S$ |
| --- | --- | --- | --- |
| $W_1$ | 0.52 | 0.40 | 0.53 |
| $W_2$ | 0.49 | 0.36 | 0.46 |
| $W_3$ | 0.34 | 0.25 | 0.41 |
| $W_4$ | 0.82 | 0.30 | 0.47 |

**Table 2.**
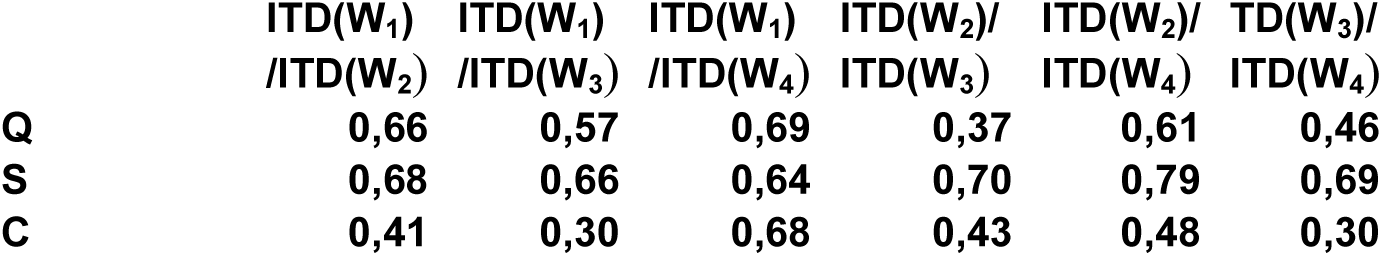
R values calculated between the different ITD(W_i_) for each studied EEG epoch.

1. input columnar activity *i*_*i*_ (*t*) during EEG epoch Q is more correlated to that occurring during S (R=.83) than that recorded during C (R=0.59) and this later is more correlated to C (R=74) than to Q (R=0.59). This may be interpreted assuming that cortical activation *i*_*i*_ (*t*) during the different studied epochs, although sharing many common CCAs, has a distinct a distinct distribution during the different phase of quiz solving;
2. each EEG component W_i_ has a distinct input cortical activation signature when compared one to each other, as well as when the different quiz solution phase is considered and
3. SLF is high for hub nodes and exponentially decreases for subordinate nodes.

All these observations reinforce the necessity to combine technologies to get better results from EEG analysis.

### 3.5 Sequential ILS location

Figure 18 shows the results of analysis of the sequential order at which CCAs are chosen as first solution of the inverse problem. In this analysis, given that the spatial location s_xyz_(t) at x,y,z coordinate is selected at time t as the best inverse problem solution, then the previous s_xyz_(t-1) and the next (succeeding) s_xyz_(t+1) best solutions are selected for study.

**Figure 18.**
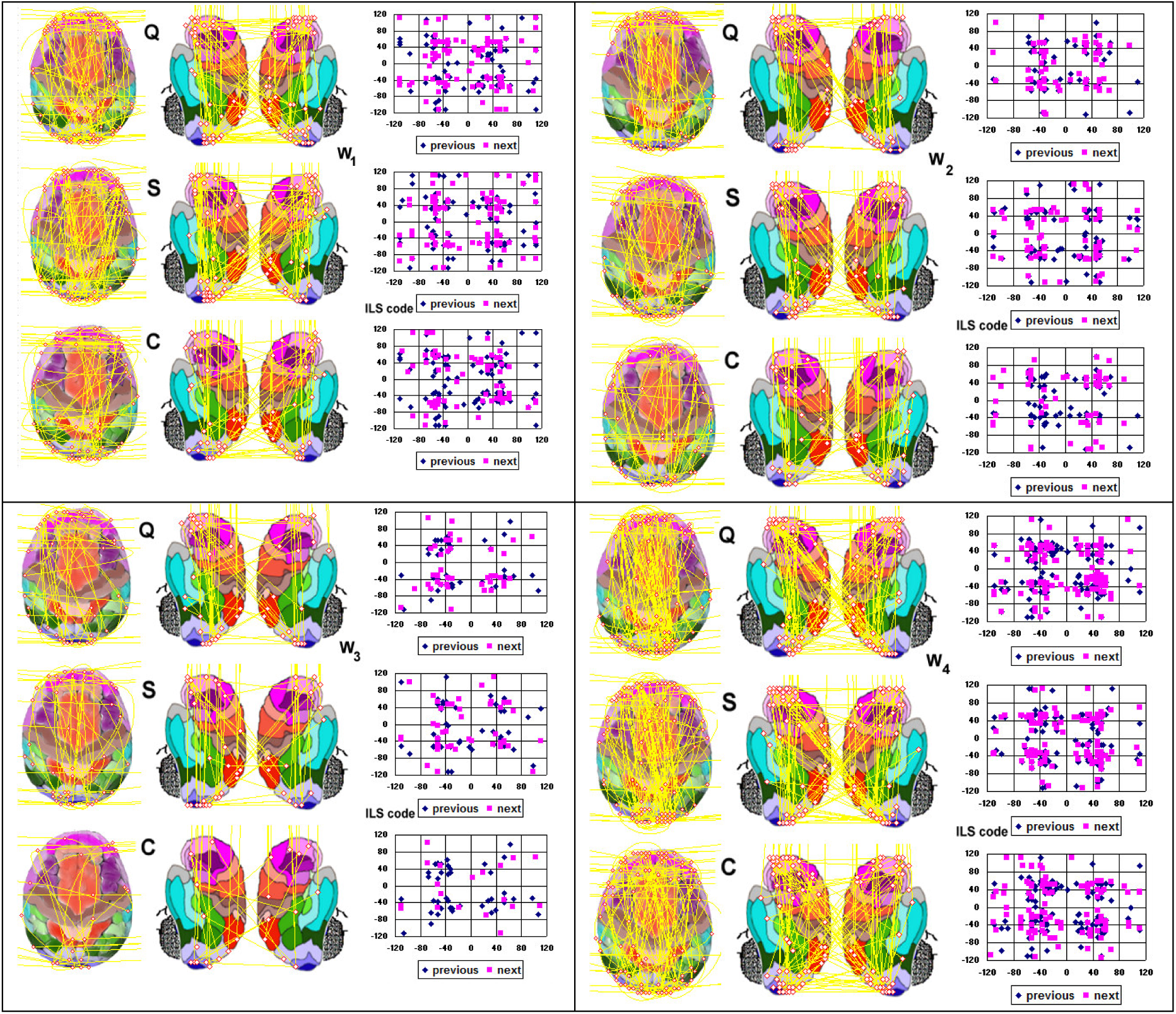
Sequential identification of ILS considered as first solution of the inverse problem. The *actua*l selected location s_xyz_(t) serves as reference to identify previous and next selected areas. Lines in the brain mappings (at left) connect these *previous* s_xyz_(t-1) and *actua*l selected location s_xyz_(t) locations. Lines have no direction so they may link previous to actual locations or vice versa. Graphics at right display s_xyz_(t-1) (blue diamond) and s_xyz_(t+1) (prink squares) locations (Y axis) referred to s_xyz_(t). Lines running out to right, left or up mean that either previous or next sources were not statistically significant.

Inspection of the brain mappings in figure 18 shows that source alternation occurs, for all studied EEG epochs, predominately between sources located at BA 10/11 and BA18/19 in the same or distinct hemispheres, as well as between BA 10/11 and BA18/19 located at both hemispheres. Alternation seems to predominate in the left hemisphere except for EEG epoch C exhibiting a right source alternation predominance.

Analysis of the graphics in figure 18 corroborates the above observations, showing four clear clusters of sources encoded in the range 30 to 60 for both hemisphere and all EEG epochs and components. These results shows that actual sources s_xyz_(t) located at BA 10/11 and BA 18/19 are preceded (s_xyz_(t-1)) and succeeded (s_xyz_(t+1)) by sources locally in the vicinities of BA10/11 or BA 18/19 or distantly at BA 10/11 and BA 18/19 (or vice versa) located at the same or opposite hemisphere. Besides these clear alternation predominance, any s_xyz_(t) are also preceded and succeeded by s_xyz_(t-1) and s_xyz_(t+1) located at many other different cortical areas. These results are confirmed by correlation analysis between W_i_ calculated ITDs shown in table 3.

**Table 3.** R values calculated between the different ITD(W_i_) for each studied EEG epoch.

|  | ITD(W<sub>1</sub>)<br>/ITD(W<sub>2</sub>) | ITD(W<sub>1</sub>)<br>/ITD(W<sub>3</sub>) | ITD(W<sub>1</sub>)<br>/ITD(W<sub>4</sub>) | ITD(W<sub>2</sub>)/<br>ITD(W<sub>3</sub>) | ITD(W<sub>2</sub>)/<br>ITD(W<sub>4</sub>) | ITD(W<sub>3</sub>)/<br>ITD(W<sub>4</sub>) |
| --- | --- | --- | --- | --- | --- | --- |
| previous |  |  |  |  |  |  |
| **Q** | **0,74** | **0,60** | **0,68** | **0,53** | **0,79** | **0,50** |
| **S** | **0,71** | **0,64** | **0,69** | **0,71** | **0,81** | **0,68** |
| **C** | **0,44** | **0,38** | **0,61** | **0,49** | **0,60** | **0,44** |
| next |  |  |  |  |  |  |
| **Q** | **0,69** | **0,61** | **0,68** | **0,50** | **0,75** | **0,52** |
| **S** | **0,69** | **0,62** | **0,69** | **0,67** | **0,80** | **0,58** |
| **C** | **0,40** | **0,34** | **0,61** | **0,48** | **0,53** | **0,42** |

Data displayed by brain mappings and graphics clearly shows that cortical activation differs for each W_i_ component and each EEG epoch Q, S and C. These results strongly support the view that cortical processing recruits neurons located at widely distributed cortical areas having a complex dynamics.

The results in this section together those about hub nodes in the previous section suggest that activity hub nodes at BA 10 and 11 alternates with that at hub nodes BA 18 and 19. A less frequent alternation involves hub CCAs located at BA 47.

### 3.6 Identifying the columnar assemblies generating oscillatory activity

LORETA solves the inverse problem of identifying the possible columnar assemblies contributing to the genesis of each of selected frequency using the calculated TVCRS (see figure 13) for a given EEG epoch. Figure 19 shows the results for this kind of analysis in the case of the EEG epochs Q, S and C. As pointed above, LORETAs allows the selection of up to 5 of the most probable solution, but it provides accurate x,y,z coordinates only for the first of then. Figure 19 display the spatial location of the first solution.

**Figure 19.**
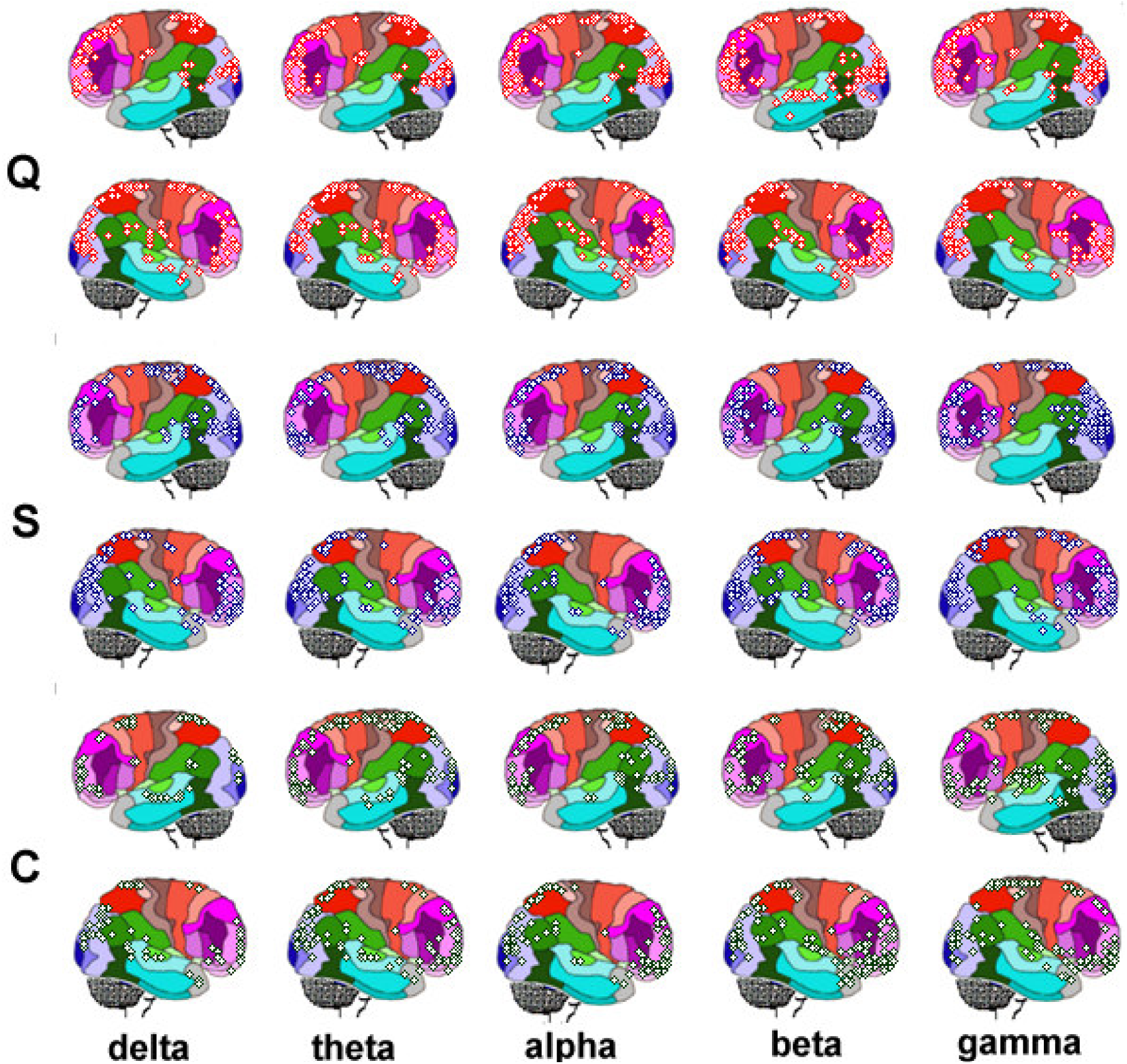
Spatial location of the Identified Loreta Sources (ILS) as possible generator of the distinct studied band frequencies. Delta - 1 to 3 Hz; Theta - 4 to 7 Hz; Alpha - 8 to 13 Hz; Gamma - 14 to 25 Hz, and Gamma - above 25 and smaller than 65 Hz.

As pointed before, selection frequency location of spatially ordered ILSs for the different EEG epochs and different EEG components may be compared using correlation analysis, because spatially ordered ILSs may be considered as spatial series. Figure 20 shows the results of this type analysis for the different EEG epochs and studied band frequencies (delta - SLF_D_; SLF_T_ - theta; alpha - SLF_A_; SLF_B_ - beta and SLF_G_ - gamma) using the same data displayed in figure 19.

**Figure 20.**
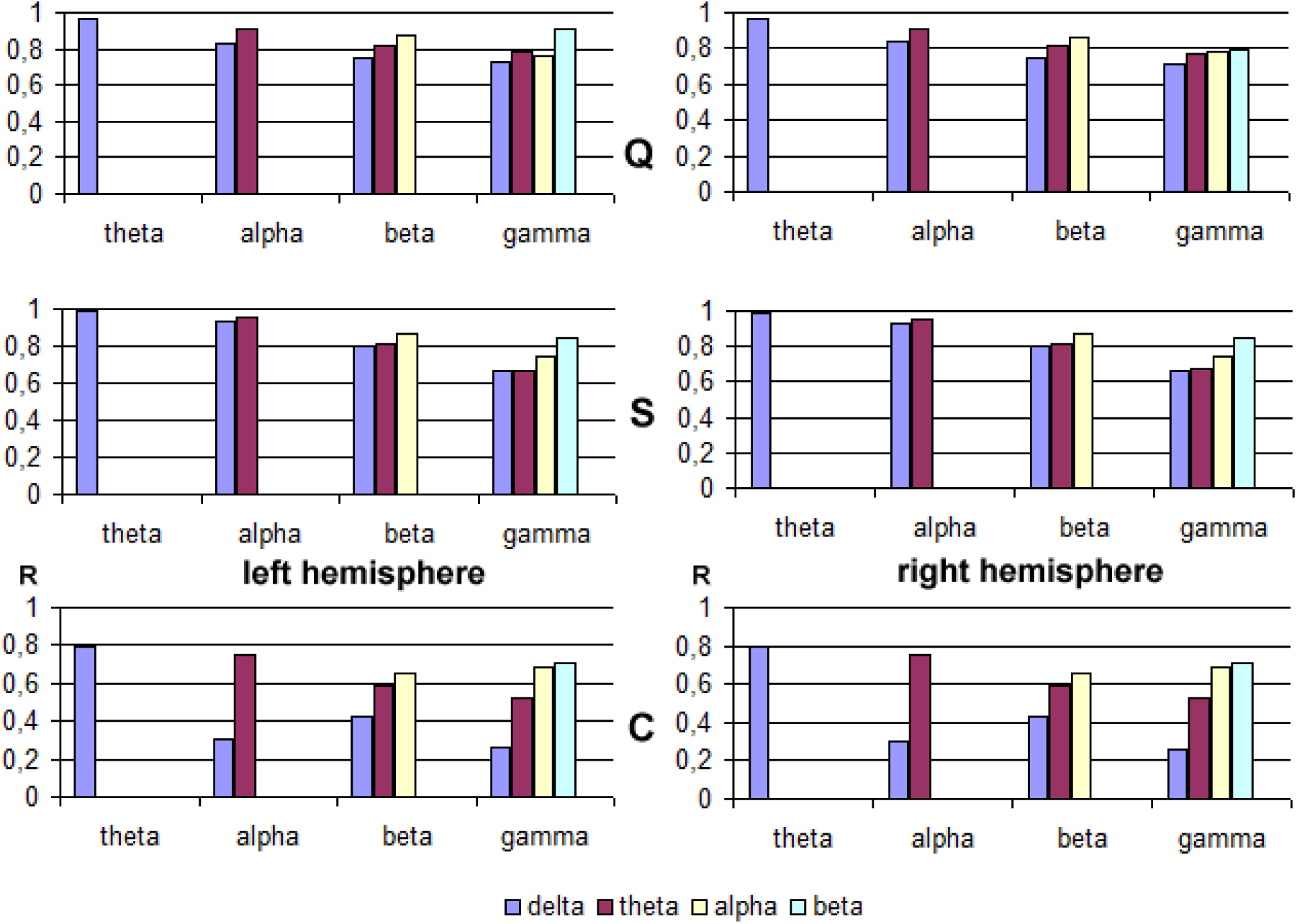
Correlation analysis of the cortical spatial location of the identified ILS shown in figure 16 taking into account the different band frequencies and each cerebral hemisphere. Graphics show the R values for the correlations specified by color column and Y axis label. For example, first blue column encodes R value for the correlation between SLF_D_ x SLF_T_; the second dark red column encodes R value for SLF_D_ x SLF_A_, etc. The last green column encodes R value for SLF_B_ x SLF_G_.

Inspection of figure 20 shows that all correlations between all frequencies are high for EEG epochs Q and S in comparison to C. Also, there is a trend to R be higher for close band frequencies (e.g., delta and theta) in comparison to R calculated for distant band frequencies (e.g., delta and gamma). These results may be interpreted by assuming that close band frequencies tend to have generators at each cortical location closer to each other in comparison to the sources for distant band frequencies

Combining information provided by figures 19 and 20, it may be considered that generators of the different band frequencies tend to, but not exclusively, occur in the same cortical neighborhood.

Figure 21 shows ILS temporal distribution for the studied band frequencies considering the different EEG components W_i_ for Q, S and C. Table 4 shows the results of the correlation analysis for the Band Source Frequency Location (BSLF) for summation of the ILS frequency calculated for each band frequency, taking into consideration the different W_i_s and each EEG epochs.

**Table 4.** R values calculated for SLF epoch relations and each W_1_.

| | $BSLF_Q/BSLF_S$ | $BSLF_Q/BSLF_C$ | $BSLF_C/BSLF_S$ |
| --- | --- | --- | --- |
| $W_1$ | 0,32 | 0,30 | 0,68 |
| $W_2$ | -0,05 | 0,08 | 0,80 |
| $W_3$ | 0,18 | 0,16 | 0,46 |
| $W_4$ | 0,25 | 0,15 | 0,77 |

**Figure 21.**
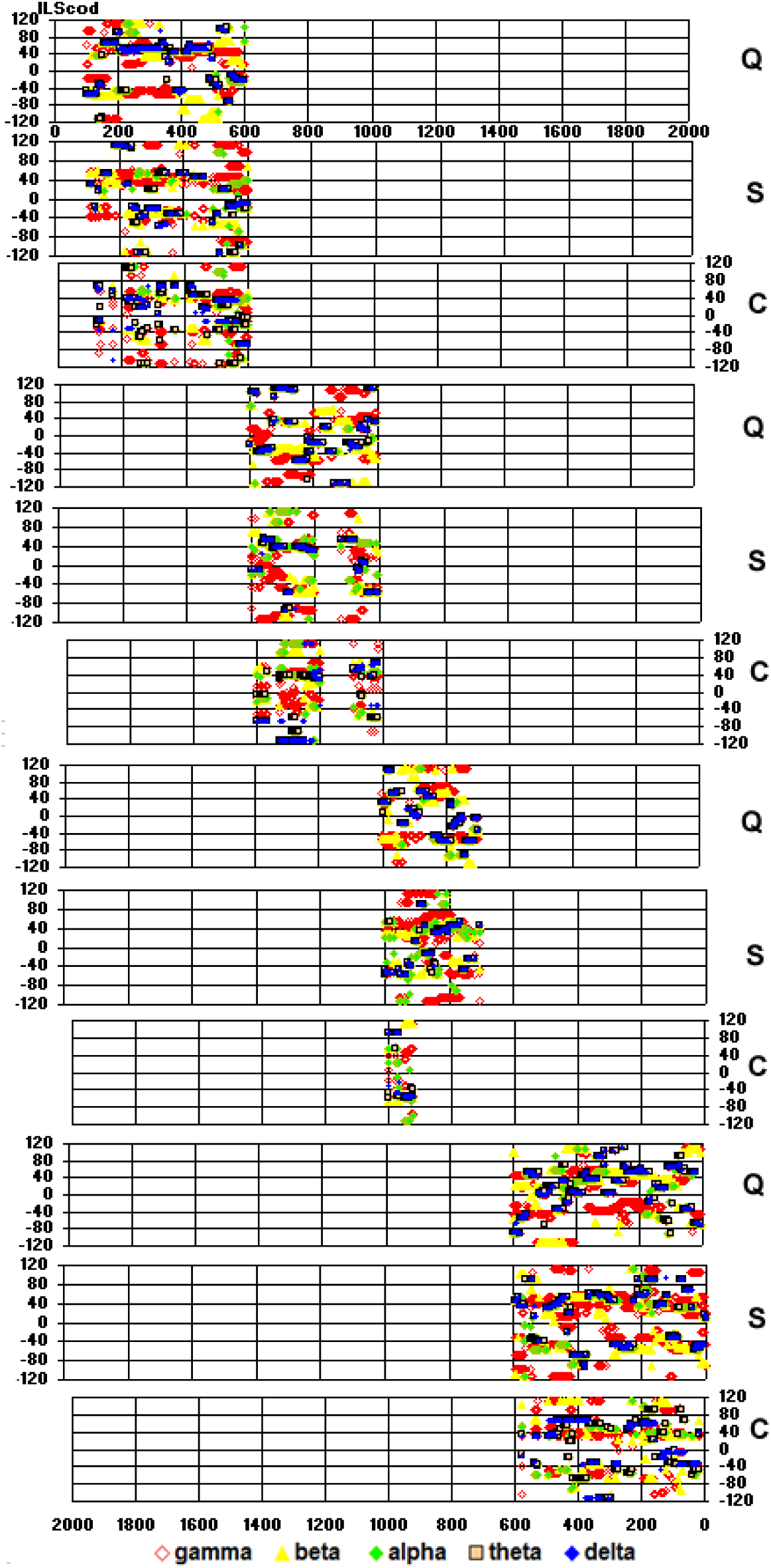
Band Frequency ILS temporal distribution associated to each EEG component W_i_ and EEG epochs Q, S, and C.

Inspection of data displayed in Figure 16 shows that identified cortical sources (ILSs) of each band frequency varied according to the studied EEG epoch Q, S and C, as well as each EEG component W_i_. Table 4 shows that these differences are smaller for comparison of source frequencies obtained for epochs S and C.

These results supports the conclusion that each EEG component W_i_ has their particular cortical oscillatory activity signature, in the same way they have their distinct input columnar activity signature, too.

Regression analysis disclosed a exponential relation between SLF and *h*(*e*_*i*_) as shown in figure 2. Hubness is high for CCA located at BA 10 or 11, BA 18 or 19 and BA 47 no matter the considered EEG epoch. CCAs located at BA 7, BA 10 and 11, BA18 and 19 and BA 47 were the most important hub nodes for all EEG epochs. BA 7 was found to be a hub in case of epochs Q and S, while BA 21 was found to be a hub in case of epoch C.

### 3.7 Correlating ILS location and cortical entrainment patterns as characterized by FA

Figure 23 shows brain mappings of all ILSs calculated for input activity *i*_*i*_ (*t*) (amplitude) and band frequencies (frequency) activity *i*_*c*_ (*t*) and of their correlation with FA patterns of cortical entrainment. Inspection of the figure shows that ILS spatial location nicely correlates with the identified entrainment patterns P_i_.

**Figure 22.**
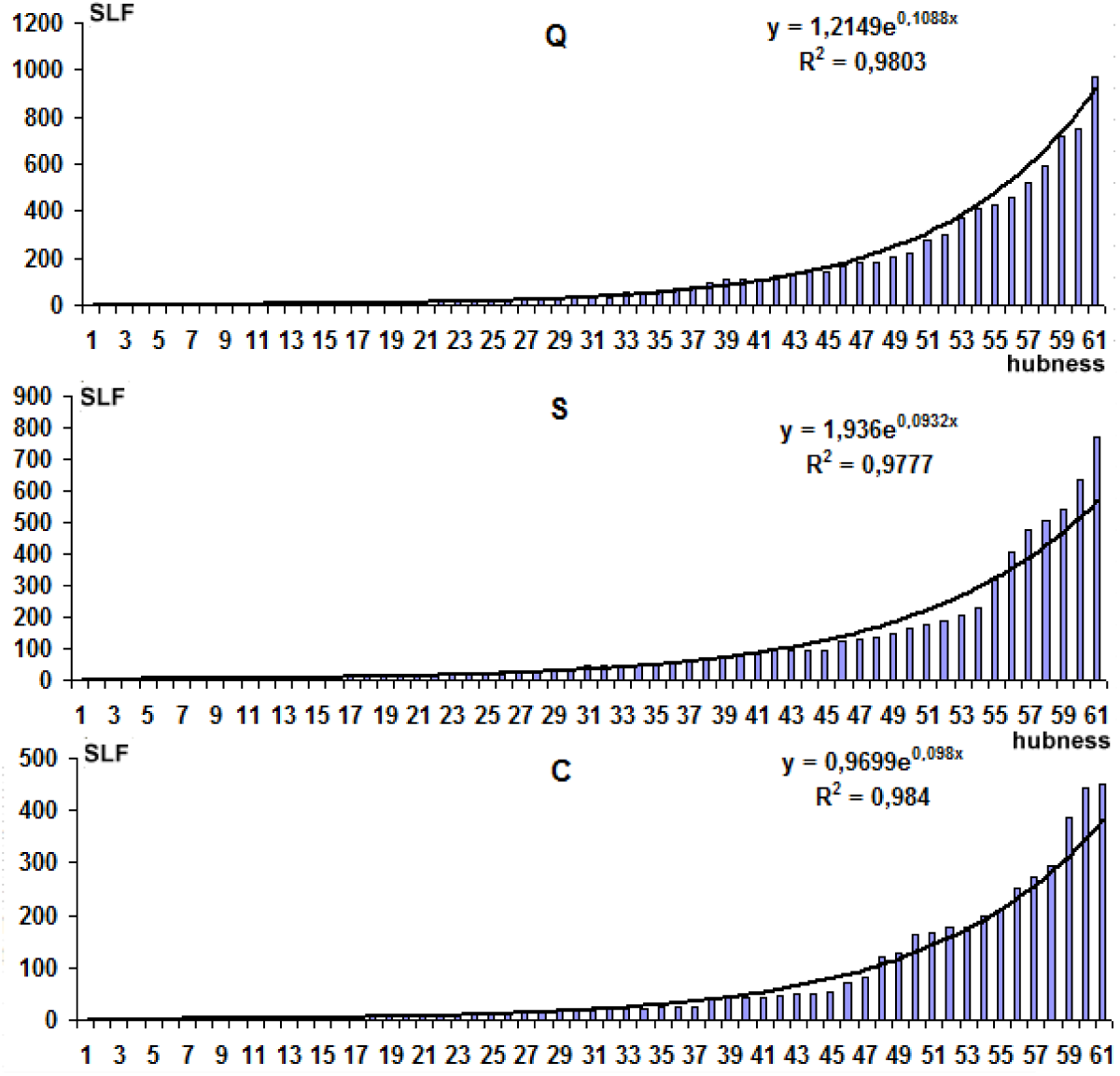
Oscillatory sources Hubness - SLF distribution follows a exponential law when is ordered in a increasing fashion and plotted according to the corresponding order index or hubness.

**Figure 23.**
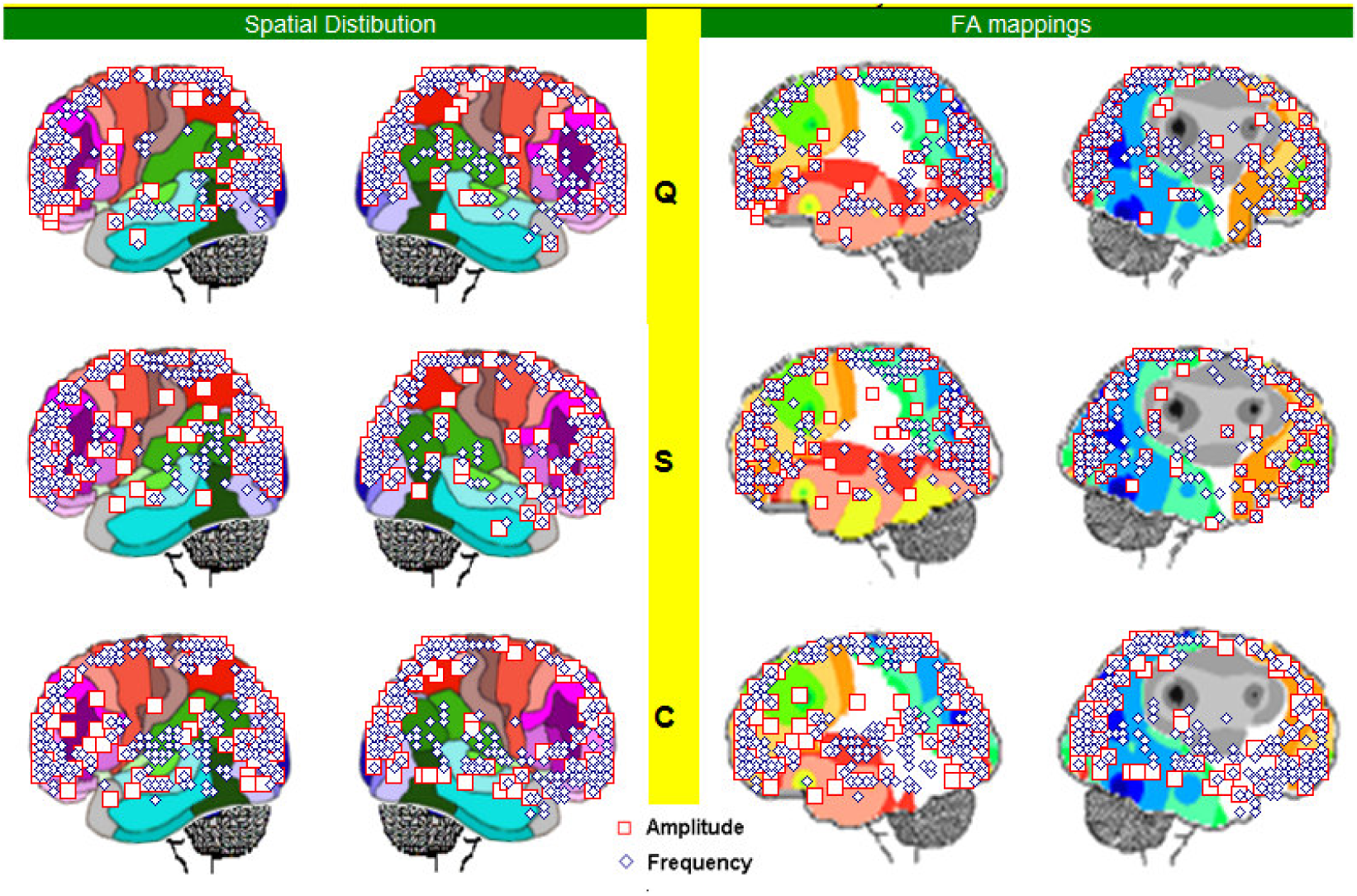
Input activity (amplitude) and band frequencies (frequency) ILSs and their correlation with FA entrainment patters for EEG epochs Q, S and C.

ILS located at BA 5 and 7 as well as BA 17, 18 and 19 are those closest to the electrodes heavily loading on FA entrainment pattern P_1_. Sources located at BA 6, 8, 9, 10 and 11 as well as at BA 44, 45 and 46 are those nearest to the electrodes loading at FA entrainment pattern P_2_. In addition, ILSs located at right BA 5, 6, 7 and 40 are closely related to the electrodes loading on Pattern P_4_. Finally, sources that might have contributed to entrainment identified by pattern P_3_ are located at left BA 21, 22, 37, 38, 39, 41, 42, 43 and 44.

If it is accepted that entrainment patterns identified by FA disclose the activity of those EEG sources that are closest to the electrodes heavily loading in these factors, then it may be proposed that distinct entrainments are established between those CCAs involved in quiz reasoning. In addition, this line of reasoning seems to confirm the proposal in section 6 that quiz understanding and solution involved the entrainment of:

1. sources located at BA 5, 7, 18 and 19 in charge of the necessary visual analysis as disclosed by association to pattern *P*_1_;
2. sources located at left BA 21, 22, 37, 39, 42, 42, 43, 44 and 45 in charge of verbal analysis as revealed by their association to pattern *P*_2_;
3. sources located at BA 6, 8, 10, 10, 11, 44, 45 and 46 in charge of organizing cortical processing as disclosed by their association to pattern *P*_3_ and
4. sources located at right BA 5, 6, 7 and 40 in charge of selecting the correct picture as quiz solution as revealed by their association to pattern *P*_4_.

## 4 Conclusion

Many authors (e.g., Ghitza, 2011, Giraud and Poeppel, 2012; Oblesser et al, 2015) remarked that the relation between the time scales present in speech and the time constants underlying neuronal cortical oscillations is a reflection of and the means by which the brain converts speech rhythms into linguistic segments. In this context, speech onset triggers cycles of neuronal encoding at embedded syllabic, phonemic and phrasal scales and speech decoding becomes a process controlled by a time-varying, hierarchical window structure synchronized with the input (Ghitza, 2011). Ghitza (2013) remarked that phonetic features (duration of 20–50ms) are associated with gamma (>40 Hz) and beta (15–30Hz) oscillations; syllables (mean duration of 250ms) with theta (4–8Hz) oscillations, and sequences of syllables and words embedded within a prosodic phrase (500–2000ms) with delta oscillations (<3Hz). In a similar line of argumentation, VanRullen (2016) proposed that perception and cognition operate periodically, as a succession of cycles mirroring the underlying brain activity oscillations and proposed that, perhaps, verbal and visual systems operate under distinct oscillatory signatures (VanRullen et al, 2014).

Rocha et al (Pereira Jr, Foz, and Rocha, 2017; Rocha et al, 2015, 2017) identified a modular ERA segmentation in many distinct cognitive tasks and, because of this, they proposed reasoning to be supported by a set of modular cycles of message exchange between local and distant sets of neurons, organized by a complex cyclic multimodal oscillatory synchronized brain activity. Here, this modular oscillatory activity was detailed and its possible cortical sources identified. Let this type of cyclic modular cortical processing supported by a multimodal oscillatory activity, be called Cortical Oscillatory Modular Processing (COMP). Each processing module involves a well structured phase locking of many distinct local and distant oscillatory circuits to support the many different computations required to solve a cognitive task. These distinct synchronizations of many different oscillatory activities creates distinct patterns of *v*(*t*) variation that were identified, here, as distinct ERA_i_ components disclosed by defined sinusoidal segmentation functions. Although these components are periodical EEG events, they are not equivalent to the traditional cortical oscillatory band frequency because they result from the synchronization of these same oscillatory activities.

REM assumes that low delta activity (< 1 Hz) adjust the COMP module size to adequately support a specific cortical processing. High delta activity (1 to 3 Hz) sets the amplitude and duration of the W_i_ oscillatory phases supported by theta activity (4 to 8 Hz). Alpha, beta and gamma activities set the complementary oscillatory processing activities v_j_. REM reconstructs *v*(*t*) from sinusoidal functions representing these ERA components. It must keep in mind, that REM does not reconstruct *v*(*t*) from any EGG decomposed oscillatory activity by any specific method, e.g. Fourier analysis. This would be a non sense redundant approach. REM reconstructs *v*(*t*) from ERA components revealed by specific sinusoidal segmentation function. Hence, it is named reversed reconstruction, following the basic notion of the Reversed Engineering.

Results of REM calculations to reconstruct Quiz ERA_i_ are shown in Figure 20. The accuracy of REM to recover *v*(*t*) may be measured by two different approaches. The first calculates the correlation between *v*(*t*) and *v*_*s*_ (*t*) displayed in the upper graphic of Figure 24 and resulted in R = 0.79, The second uses Linear multiple regression analysis taking the sinusoidal functions as dependent variables and *v*(*t*) / *s*(0.5) as the dependent variable. Regression analysis uses *v*(*t*) / *s*(0.5) and not *v*(*t*) as dependent variable because of the modulator role played by low delta activity discussed above (see equations in Figure 20). Table 4 shows the results of this approach and they indicate that REM accounts for 75% of Quiz recorded *v*(*t*). In addition, this analysis shows that all *s (h)* are real contributors to *v*(*t*) because the angular coefficients **β** for these functions are statistically. different from 0 at the level of p<0.01.

**Table 4.** COMP Reverse Engineering Model - regression data.

**F(7,1527)=163.48 p<0.0000 Std.Error of estimate: .47635**
| | $\beta$ | St. Err.<br>SD $\beta$ | t(2552) | p-level |
| --- | --- | --- | --- | --- |
| Intercept | 100.3566 | 39.05818 | 9.128826 | 2.13E-19 |
| S(0.5) | 100.3566 | 0.039058 | 9.128826 | 2.13E-19 |
| S(1) | 74.4259 | 12.10522 | 6.148247 | 9.98E-10 |
| S(2) | -34.9535 | 2.447854 | -14.2792 | 1.64E-43 |
| S(5) | -92.0562 | 5.236421 | -17.58 | 0 |
| S(14) | -18.1091 | 0.900163 | -20.1175 | 0 |
| S(30) | 113.6285 | 34.50394 | 3.293203 | 0.001013 |

**Figure 24.**
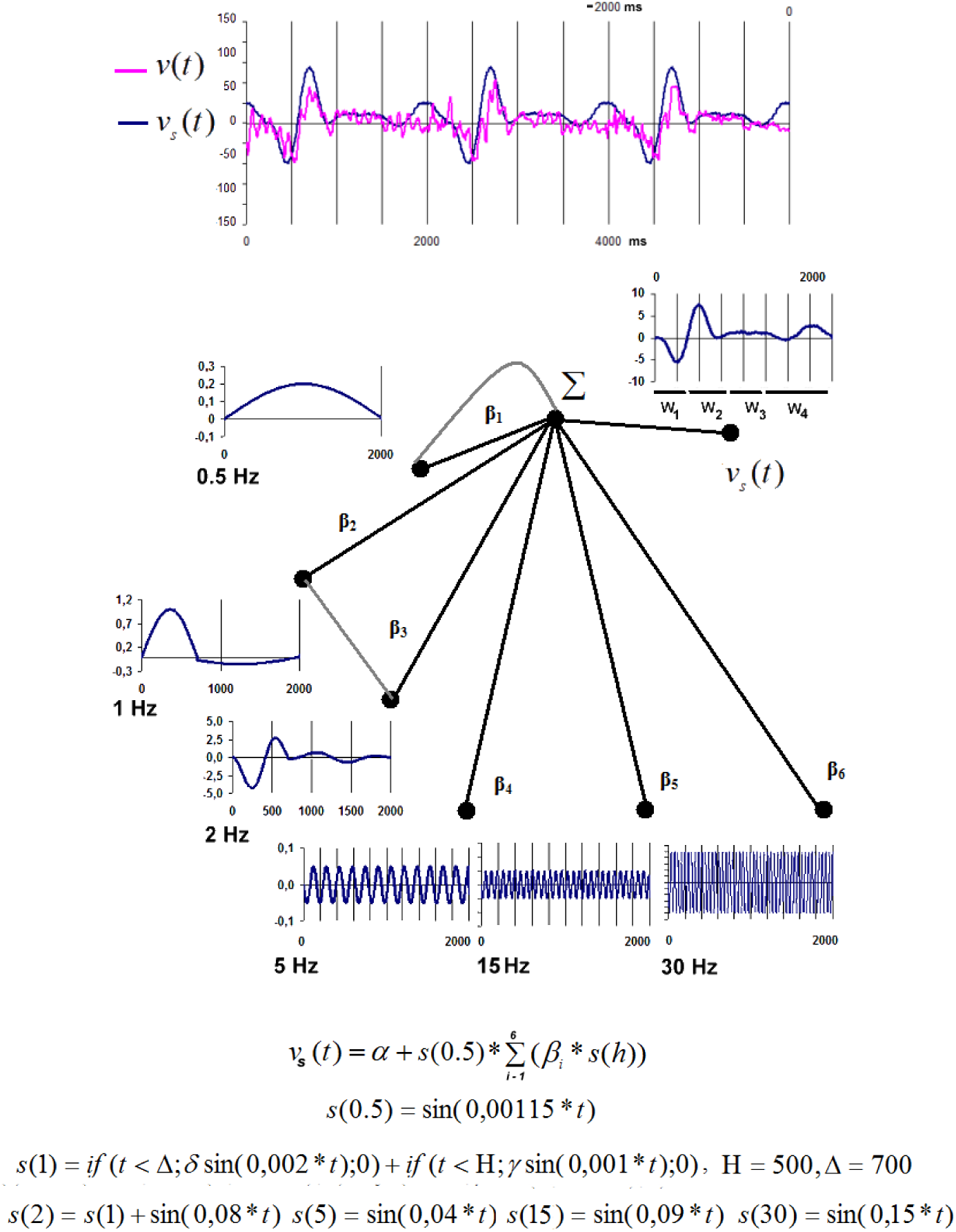
The regression engineering model (REM) The simulated *v*_*s*_ (*t*) (the dependent variable) is calculated from a set of sine functions (independent variables) s(0.5) to s(30) as 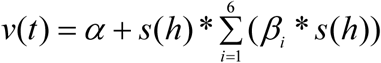 *-* The inserted graph shows the structure of this model. Its terminal nodes represents each one of the dependent variables and the arcs shows how they contribute to *v*_*r*_ (*t*), that is represented at the graph root node. Dependences of s(2) on s(1) and of *v*_*s*_(*t*) on 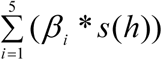 calculation are shown in gray and all other dependences are show in black. - Upper graph shows the real averaged *v*(*t*) and the recovered *v*_*s*_ (*t*) Linear regression analysis may be used to check the plausibility of the proposed model COMP, using sinusoidal functions as dependent variables to reconstruct *v*(*t*) as the independent variable. Here, this reverse *v*_*s*_ (*t*) reconstitution is called Reversed Engineering (Regression) Modeling (REM). If COMP is supported by a structured modular set of computations carried out by many distinct oscillatory neural circuits, then a regressed *v*_*r*_ (*t*) computed from a set of sinusoidal functions simulating these distinct oscillatory activities, must be well correlated to the recorded *v*(*t*).

Burgess (2012) proposed that ERA waves are generated by synchronization of various oscillatory activities in many distinct cortical areas. A view supported also by Klimesch et al (2007). The similarity of both the calculated ERA_i_ and its components may be understood by assuming that this oscillatory phase locking within and between oscillatory band frequencies determines the actual values of recorded *v*(*t*) and the corresponding ERA_i_ components.. Oscillatory activity is generated by many distinct sets of neurons widely distributed over the cortex. Many of these sources are common for the different ERA_i_ and their components, as well as for distinct studied oscillatory band frequencies. In addition, other sources are specific to each COMP cycle. As a consequence, the shape similarities may be accounted by phase locking of similar oscillatory activities while at the same time shape differences may be accounted by variation of both oscillatory activity and source location. REM seems to reinforce these propositions by showing that common ERA features are simulated with the same set of oscillatory functions mimicking ERA_i_ components..

Here, oscillatory phase locking is assumed to be promoted by the entrainment supported by the scale free structure of the COMP neural circuits. Low delta (e.g, 0.5 Hz component) COMP modulation is crucial to the concept of modular processing because 1) it constrains phase locking of all other oscillatory activities to a defined period of time, therefore determining the size module. High delta (e.g., 1 Hz component) activity controls W_i_ variability within each COMP cycle, and consequently, the duration of the computations supported by components W_i_ and w_j_ created by theta and alpha activities, respectively. In the same line of reasoning, beta and gamma activities impose the dynamics of other REM independent variables. In this model, low oscillatory activity imposes a top down control upon the bottom up entrainment promoted by input activity as discussed by Zoefel1 and VanRullen (2016). Lakatos et al (2008, 2010) showed that when attended stimuli are in a rhythmic stream, delta-band oscillations in the primary visual cortex entrain to the rhythm of the stream, resulting in increased response gain for task-relevant events and decreased reaction times. Because of hierarchical cross-frequency coupling, delta phase also determines momentary power in higher-frequency activity.

The above assumptions are also in line with the observations of von Stein A and Sarnthein (2000), showing that while local gamma synchronization is involved in visual processing, beta synchronization occurs between temporal and parietal neighboring areas during multimodal semantic processing and alpha and theta synchronization are observed in fronto-parietal circuits supporting working memory and mental imagery. Based on these results, the authors suggested a relationship between the extent of functional integration and the synchronization-frequency. They also assumed that low frequency ranges seem specifically involved in processing of internal mental context, i.e. for top-down processing.

All these results refute any reductionist approach of the cortical function trying to assign complex cognitive functions to specific sets of neurons and to specific time event. Instead of this, they show that the available tools for EEG analysis are very useful to investigate cortical dynamics of human reasoning if cerebral processing is assumed to recruit a large number of cells, both neurons and astrocytes, located at innumerous cortical areas.

A key issue in a distributed processing supported by a large number of agents like the brain, is the activity entrainment of agents recruited to solve a given task. Columnar structure and function supporting oscillatory activity favors both local and distant entrainment *E*(*i*) of the CCA activated during any kind of reasoning. Entrainment is enforced by columnar input activity*i*_*i*_ (*t*) derived from sensorial and motor systems; thalamus and other columnar assemblies. Tools for EEG analysis described and discussed in the present paper are very adequate to investigate and characterize *E*(*i*).

Results discussed here and elsewhere support the proposition that:

1. cortical processing is supported by modular sets of computations (COMP) of around 2 seconds duration each;
2. involving many distinct oscillatory phases that are easily identified by ERA analysis;
3. that are generated by different sets of columnar assemblies and
4. phase locked by entrainment *E*(*i*) of distinct scale free neural circuits involved in handling a given cognitive function.

In this line of reasoning, it may be concluded that EEG must be the first tool of choice to study human cortical cognitive activity as proposed by Berger (2009) almost a century ago, due to its very high temporal discrimination of the distributed cortical processing dynamics. But this preference is only supported by a combined use of the different available tools for analysis of the distinct types of information encoded in the recorded *v*(*t*).

## Footnotes

1 www.uzh.ch/keyinst/NewLORETA/Software/Software.html

